# Ratiometric transcriptional activation by protein degradation

**DOI:** 10.64898/2026.05.16.725679

**Authors:** Melissa A. Gray, Katelyn L. Randal, Jennifer A. Co, Michelle T. Tang, Athena Z. Xue, Sophia W. Chen, Hlib Razumkov, Qusay Q. Omran, David. E. Solow-Cordero, Jeonghye Yu, Stephanie A. Robinson, Cara A. Starnbach, Nathanael S. Gray, Steven M. Corsello, Steven M. Banik

## Abstract

Cells can respond to alterations in the abundances of specific proteins through transcriptional outputs. Synthetic approaches inspired by native post-transcriptional circuits that convert protein abundance changes into programmable gene expression would be transformative. Here, we discover and describe design principles that effectively convert protein degradation into transcriptional outputs in live cells. We define ratiometric transcriptional activation, where control over the ratio between a transcriptional inhibitor-protein of interest fusion and transcription factor enables detection of abundance changes with high sensitivity at scale. We show that ratiometric transcriptional activation can be implemented in single cells using triply orthogonal circuits or in multicellular pools, operating independently of mechanism of protein downregulation and enabling simultaneous detection of multiple protein downregulation events through outputs such as cell survival, fluorescent protein expression, or barcode sequencing. These circuits can be applied to oncogenic targets and enable discovery of new molecular glue degraders.

## Introduction

Protein degradation is a defining feature of cellular homeostasis.^1^ Cells dynamically respond to stimuli by rapidly manipulating both global^2^ and specific^3^ protein levels. Biological circuits constantly sense the presence or absence of proteins and convert this information into coordinated responses. Altered proteostasis is a marker of disease states including cancer^4,5^ and neurodegenerative disease,^6^ where post-translational changes in protein abundance can lead to unregulated growth or neurological decay, respectively. Global alterations in proteostasis at the post-translational level through proteasome and lysosome inhibition have led to key therapeutics for multiple indications.^7,8^ The targeted degradation of proteins of interest has provided unprecedented precision in manipulating post-translational biology.^9,10^ Therapeutically, the concept of protein degradation presents an alternative mechanism to impact difficult to drug proteins beyond direct target inhibition.^11–14^

Detection of protein levels is essential for both therapeutic development and fundamental biology. Current methods for identifying conditional protein loss include proteomics,^15^ immunoblotting, or monitoring signals from a fused fluorescent^16^ or bioluminescent reporter.^17^ All methods tie direct loss of the target protein to loss-of-signal (e.g. down-assays). This prevents signal amplification, limits the types of observable signals, and provides few options for increasing sensitivity beyond target protein overexpression. An alternative to sensing direct loss-of-signal is to tie protein degradation to cell survival;^18^ however, the amount of protein degradation needed to observe survival will vary based on target protein expression level and dynamics. A method which releases the constraints on protein abundance, bypasses the direct correlation of protein depletion to loss-of-signal, and provides a tunable and multiplexable signal upon target protein downregulation could enable sensitive detection of protein regulators and programmable biological outputs at scale.

Programmable genetic circuits can interface with native biological systems to generate reporters of cellular activity. Circuits have been developed that sense proteolytic activity,^19,20^ extracellular environments,^21^ cell-cell interactions,^22^ and post-translational modifications.^23^ However, no current genetic circuit transforms protein degradation into flexible cellular outputs. Cells maintain proteins at specific ratios to regulate transcription; for example, NF-κB is inhibited by the IκB family of proteins; upon IκB degradation, transcription is rapidly induced by free NF-κB (Figure 1A).^24^ A synthetic approach using a functional internal standard that modulates transcription would allow for amplified detection of protein downregulation and tie loss of a protein to a gain of programmable signal, a concept we term “ratiometric transcriptional activation” (RTA) (Figure 1B). Here, we program and characterize design principles of genetic circuits to provide an amplified gain-of-signal upon downregulation of a target protein and deploy them for small molecule discovery.

**Figure 1.**
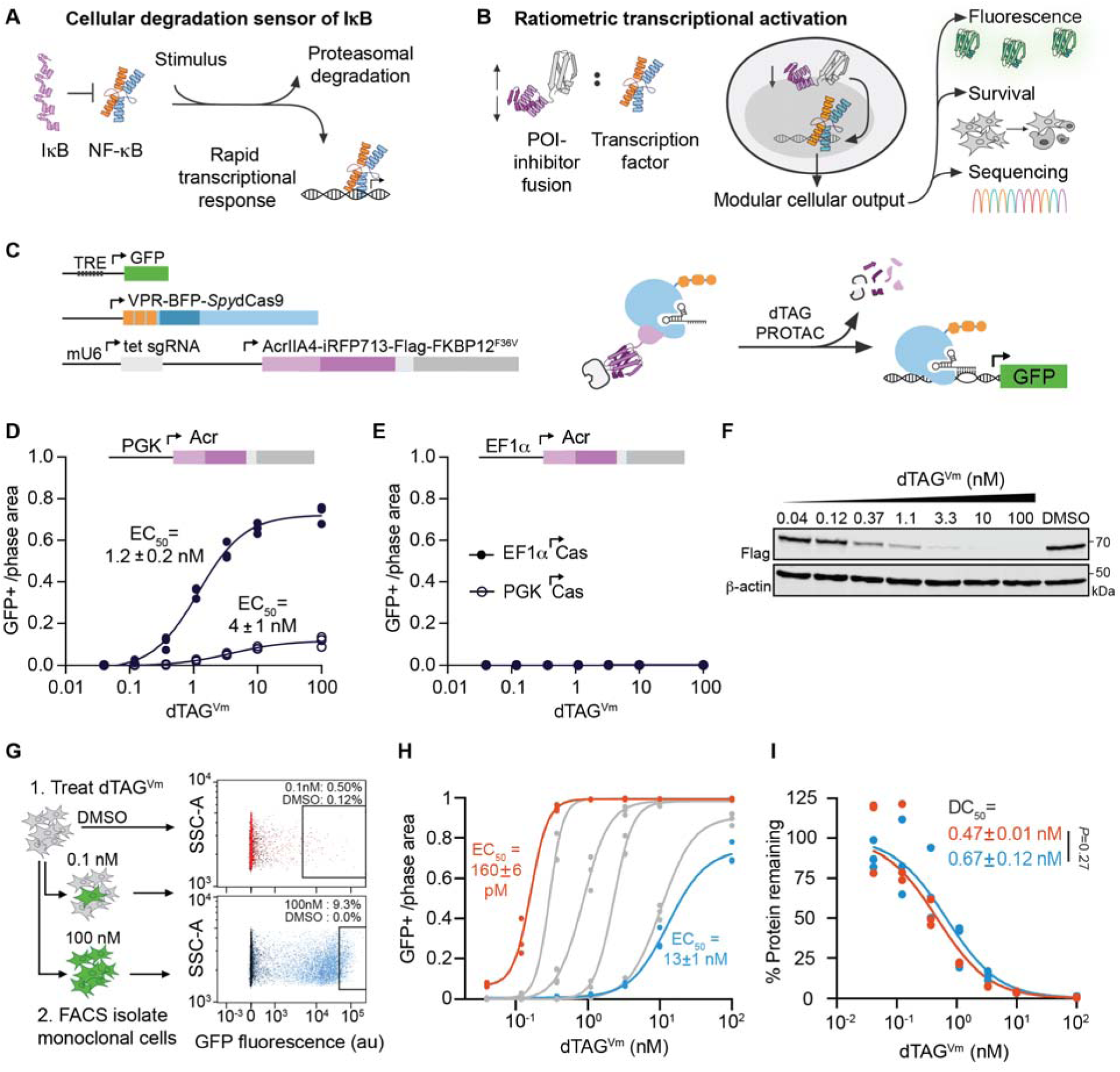
Ratiometric transcriptional activation converts tunable levels of protein degradation into amplified outputs. (**A**) Cells tune transcriptional activation by perturbing protein ratios. The ratio of IκB to NF-κB transcription factor inhibits NF-κB, which is released upon stimulus and degradation of IκB. (**B**) Illustration of the concept of ratiometric transcriptional activation (RTA), in which a protein-of-interest (POI) is fused to a transcription factor inhibitor. Decreased concentration of inhibitor relative to the internal standard transcription factor promotes gene expression. (**C**) A model system that detects FKBP12^F36V^ protein degradation by its fusion to AcrIIA4 (Acr) to inhibit VPR-*Spy*dCas9 (dCasA) activity. (**D-E**) Live cell fluorescence microscopy quantification of GFP+ area normalized to total cell area of polyclonal circuit-containing cells with (D) PGK- or (E) EF1α-promoted dCasA treated with increasing concentrations of dTAG^Vm^ (n=3). (**F**) Representative immunoblot of PGK-promoted Acr-iRFP713-Flag-FKBP12^F36V^ after 24 hours of dTAG^Vm^ treatment. (**G**) GFP+ monoclonal cell lines were sorted from the polyclonal parent line after treatment with low (0.1 nM) or high (100 nM) dTAG^Vm^ for 24 hours. (**H**) Comparison of six monoclonal cell lines GFP+ area normalized to total cell area after 24 hours treatment with increasing dTAG^Vm^ concentration analyzed by live cell fluorescence microscopy (n=3). (**I**) Protein degradation as a function of dTAG^Vm^ concentration in sensitive (red) and insensitive (blue) monoclonal cell lines by quantified immunoblot normalized to DMSO for each cell line (n=3, *P* value: two-tailed t test comparing DC_50_ values). EC_50_ and DC_50_ values in this figure are calculated from four-parameter logistic equation curve fits.

### Design of a synthetic ratiometric transcriptional activation circuit

To design a sensitive protein downregulation responsive circuit, we reasoned that by fusing an inhibitor of a transcription factor to our protein-of-interest (POI), POI loss would be benchmarked to transcription factor levels in cells, leading to sensitive and amplified readouts (Figure 1B). The Cas protein toolbox provides programmable transcriptional regulators that are orthogonal to native cellular machinery. Anti-CRISPR (Acr) proteins inhibit Cas protein binding to DNA^25^ and have been used to control Cas-based genome editing, transcriptional activation and repression ^26^. The characterized Acr family member, AcrIIA4, a 10 kDa protein derived from *L. monocytogenes* prophage, binds to guide RNA-loaded *S. pyogenes* Cas9 (K_D_ ∼ 0.6 nM) to inhibit DNA engagement.^27^ The crystal structure of the AcrIIA4-*Spy*Cas9 complex reveals solvent exposed Acr termini potentially amenable to fusion with protein targets (Figure S1a).

As an initial demonstration of ratiometric transcriptional activation, we fused a direct reporter, iRFP713, as well as an FKBP12^F36V^ protein, for which small molecule degraders have been developed (dTAGs)^9,28^ to the C-terminus of AcrIIA4 (Figure 1C). These constructs were stably integrated along with a targeting guide RNA into polyclonal cells containing *Spy*dCas9-based transcriptional activator (dCasA) and a TRE (7x tetO)-promoted GFP reporter gene. Stable integration was chosen over transient plasmid expression to prevent saturation of cellular machinery needed to degrade target proteins. To assess the contribution of relative expression levels of inhibitor and transcription factor to circuit behavior, we generated a panel of four polyclonal stable circuits with dCasA and Acr expression each driven by canonically weak (PGK) or strong (EF1α) promoters.^29^ All cell lines had low background indicating Acr fully inhibits dCasA-mediated GFP transcription at baseline (Figures 1D and E). Treatment of PGK-promoted Acr proteins with different concentrations of a dTAG analog (dTAG^Vm^, Figure S1b) resulted in GFP production with an EC_50_ of 1.2 nM and 4 nM by live cell fluorescent microscopy when dCasA was expressed with a strong or weak promoter (Figure 1D), respectively, and corresponding degradation of AcrIIA4-iRFP713-Flag-FKBP12^F36V^ was observed by immunoblot (Figure 1F). Acr expression under the EF1α promoter did not result in GFP signal induction upon dTAG^Vm^ treatment (Figure 1E); corresponding iRFP713 analysis by flow cytometry revealed dTAG^Vm^ treated Acr levels were still higher than steady-state PGK-promoted Acr levels (Figure S1c). Circuits with FKBP12^F36V^-iRFP713 fused to the N-terminus of AcrIIA4 resulted in high (55% GFP+) background fluorescence levels by flow cytometry (Figure S1d). Treatment with 1 μM of dTAG^Vm^ increased the GFP-on population to 90%, indicating substantial activation of transcription upon loss of target protein. Thus, Acr operates as a Cas attenuator when fused at either terminus and its abundance is a key factor for RTA protein sensing.

Fluorescence-activated cell sorting (FACS) was performed to select sensitive cell lines by treating polyclonal populations with low amounts of dTAG^Vm^ (0.1 nM) and isolating the top 0.5% of GFP-expressing cells (Figure 1G). We also isolated GFP-expressing cells derived from treatment with 100 nM, 10 nM, and 1 nM dTAG^Vm^. From these pools, we selected 12 monoclonal cell lines exhibiting ∼2 orders of magnitude range of EC_50_ values for GFP production (from 160 pM to 13 nM) (Figures 1H and S1e). By triplicate immunoblot and measurement of iRFP713 degradation by flow cytometry, the most and least sensitive monoclonal lines did not have a significant difference in DC_50_ (Figures 1I and S1f). At 40 pM dTAG^Vm^ treatment, our circuit turn-on was detectable beyond the limits of triplicate immunoblot (Figure S1g). We examined the AcrIIA4-iRFP713-FKBP12^F36V^ expression levels across 12 monoclonal cell lines and compared them to the degree of GFP production upon treatment with 1 μM dTAG^Vm^. Direct detection of iRFP713 fluorescence linearly correlated with iRFP713 signal loss upon degradation (Figure S1h). In contrast, the initial expression levels of AcrIIA4-iRFP713- FKBP12^F36V^ were decoupled from the degree of GFP activation upon treatment (Figure S1h). Together, these data demonstrate that sensitive detection cells can be selected for both low and high amounts of degradation, and that these synthetic circuits use amplified signal to report on degradation regimes inaccessible to immunoblots.

### Generation of a quantitative model to understand ratiometric transcriptional activation

To support the RTA principle, we developed a simplified ordinary differential equations (ODE)-based model that captures the anticipated core dynamics of the system, including synthesis, binding interactions, and degradation of each species. The model predicts that Acr/dCasA ratios define the sensitivity of the cellular circuits under accelerated degradation conditions (equivalent to increasing dTAG doses in a cell) (Figure 2A). Circuits with [Acr] < [dCasA] had high background and did not appreciably exhibit a change in GFP (ΔGFP) upon simulated degradation (Figure 2B). In each modeled configuration, DC_50_ values for Acr degradation were unchanged, yet EC_50_ for GFP signal varied depending on component ratios (Figure 2B), recapitulating our findings with sorted monoclonals (Figure 1G). The model predicts that increasing Acr/dCasA levels exhibit a linear range directly correlated to EC_50_ with deviation from linearity at low ΔGFP. BFP signal from VPR-BFP-*Spy*dCas9 was too low to reliably detect by flow cytometry (Figure S2a). Instead, we quantified dCasA and Acr transcript levels in monoclonal cells with different EC_50_ values by qPCR (Figure 1G), and plotted dCasA (Figure S2b), Acr (Figure S2c), and Acr/dCasA transcript levels (Figure 2C) as a function of experimental circuit EC_50_. We found that Acr/dCasA ratio was the superior predictor of cell behavior compared to Acr or dCasA levels alone. Transcript levels predicted relative protein levels well (Figure S2d). The most sensitive cells had an Acr/dCasA ratio below 1, though slightly higher Acr/dCasA ratios are likely at the protein level since all cells had low transcriptional background. Taken together, these data suggest that the ratio of POI-AcrIIA4 to *Spy*dCas9 controls transcription, defines the sensitivity of degradation detection (Figure 2D), and establishes the concept of RTA.

**Figure 2.**
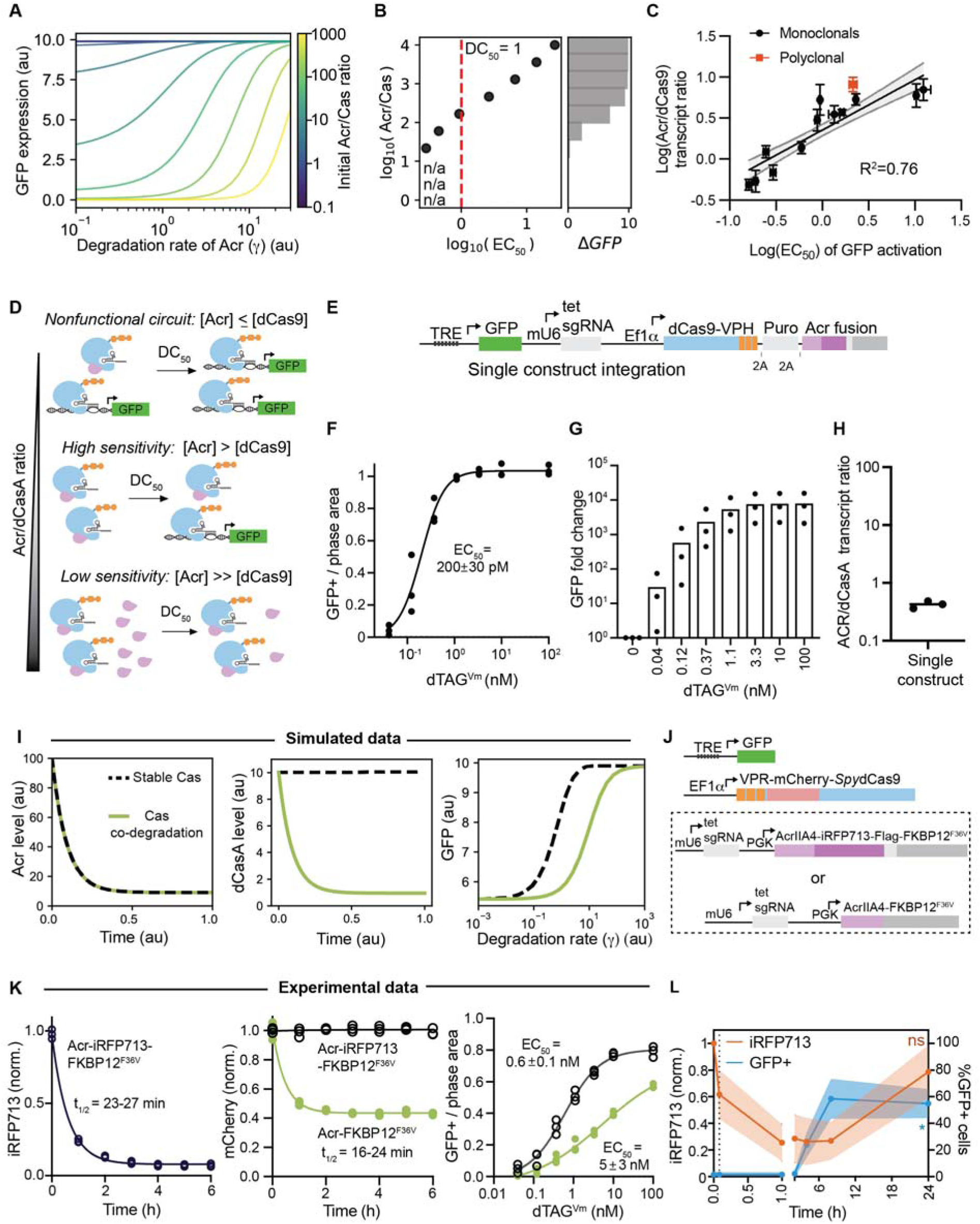
Experimental and mathematical characterization reveal design principles of RTA circuits. (**A**) A reduced-order ODE model was used to solve for steady-state GFP levels as a function of increasing Acr degradation across various starting Acr/dCasA ratios (colorbar). EC_50_ values from the simulated degradation curves are plotted in (**B**) along with the associated Acr/Cas starting ratios and total GFP increase upon maximum Acr degradation (ΔGFP). The DC_50_ value for Acr degradation across all Acr/Cas ratios is shown as red dashes. (**C**) Relative transcript Acr/dCas9 ratio measured by qPCR for 12 monoclonal cell lines with varying EC_50_ values for GFP activation. Mean ± SD and linear fit with R^2^ reported. (**D**) Summary of the proposed principle that sensitivity of circuit activation is determined by the Acr/dCas9 ratio. (**E**) An all-in-one construct stably integrated into cells contains GFP reporter, guide RNA, and dCas9A, puromycin resistance gene, and Acr expressed under a single promoter and separated by self-terminating (2A) peptides. Live cell fluorescence microscopy quantification of a polyclonal population stably expressing the all-in-one construct treated with increasing concentrations of dTAG^Vm^ (n=3), reported as (**F)** GFP+ area normalized to total cell area, and (**G**) GFP fold change normalized to DMSO controls. (**H**) Relative transcript Acr/dCas9A ratio measured by qPCR for the single construct integration cell line. (**I**) Comparison of an ODE model that incorporates co-degradation of dCas9A from the Cas:Acr complex (green line), to one that does not include Cas degradation induced by treatment (black dashed line). Plotting Acr level (left), and Cas level (middle) as a function of time after inducing substantial Acr degradation rate, and plotting GFP formation (right) as a function of increasing Acr degradation rate. (**J**) Two circuit lines were developed with either Acr-iRFP-FLAG-FKBP12^F36V^ or Acr-FKBP12^F36V^ integrated into VPR-mCherry-*Spy*dCas9 cells. (**K**) Experimental results of the lines from (J), plotting Acr (iRFP713, left) and Cas (mCherry, middle) normalized to average MFI of t=0 (untreated) samples by flow cytometry over the course of 6 hours, as well as GFP+ area over total cell area (right) by live cell fluorescence microscopy at 24 hours (n=3). (**L**) Single construct polyclonal cells were treated with 100 nM dTAG^Vm^ degrader followed by washing and replating of cells. Flow cytometry was performed at timepoints up to 24 hours to detect percentage of GFP+ cells (raw data) and Acr-iRFP713 levels (normalized to t=0 untreated control).

To simplify the implementation of RTA, we developed an expression construct requiring only a single transposase-mediated incorporation. This construct expresses all circuit components separated by the self-terminating T2A peptide to produce three proteins from the same promoter (Figure 2E), placing Acr last in the sequence to reduce Acr/dCasA levels.^30^ Upon incorporation, polyclonal cells demonstrated sensitive turn-on by live cell microscopy measuring GFP+/phase area (EC_50_ = 200 pM, Figure 2F), and by GFP fold change (Figure 2G). Quantification of relative transcript levels in the polyclonal population by qPCR confirmed an Acr/dCasA transcript ratio <1 (Figure 2H). To assess the importance of TRE promoter binding sites, we developed single expression vector circuits with 7x, 5x, 3x, and 1x sgRNA binding sites (Figure S2e), and found that sensitivity increased with binding sites. While the all-in-one design offers precise control and simplified engineering, an expanded range of component ratios is more easily accessed from a multiple construct-integration approach.

In principle, induced degradation of an Acr-fusion could result in collateral degradation of bound dCasA. We simulated dCasA co-degradation with bound Acr and observed less sensitive (higher EC_50_) circuits. However, as Acr/dCasA ratios also decrease upon degrader treatment, circuits were predicted to retain function (Figure 2I). To test this experimentally, we utilized a VPR-mCherry-dCas9 construct with detectable dCasA fluorescence, and generated two circuits, with either Acr fused directly to FKBP12^F36V^ or with an iRFP spacer between (Figure 2J). Only the Acr-FKBP12^F36V^ direct fusions showed induced degradation of dCasA upon dTAG^Vm^ treatment (Figure 2K), resulting in a reduced steady state dCasA level and a corresponding increase in transcription EC_50_. Thus, even with collateral dCasA degradation, RTA circuits remain amplified detectors of protein degradation.

Due to transcriptional turn-on and GFP stability, RTA should retain memory of protein degradation by providing a record of temporally restricted degradation events. To demonstrate recording of a transient degradation window, polyclonal single-construct cells were treated with 100 nM dTAG^Vm^ for 5 minutes followed by extensive washing and replating of the cells. After 24 hours, Acr-iRFP713-FKBP12^F36V^ recovered to a level that was similar to baseline, and robust levels of GFP were detected (Figure 2L). A similar 5 h washout experiment demonstrated GFP activation 36 h after detectable protein recovery by immunoblot (Figure S2f).

### Circuit activation with diverse degradation targets, mechanisms, and modalities

To establish generality of detection across various proteins-of-interest, we fused targets to AcrIIA4-FKBP12^F36V^ (Figure 3A). The inclusion of the FKBP12^F36V^ component enables dTAG^Vm^ treatment as a universal positive control for degradation and FACS. Our initial target protein list had well-characterized small molecule bifunctional degraders.^31^ Targets were also prioritized based on their varied subcellular localization, biological functions, and native proteostasis mechanisms. First, we generated polyclonal and monoclonal cell lines containing SMARCA4 (BRG1) (Figure S3a) and PBRM1 (Figure S3b) and observed robust signal activation upon treatment with dTAG^Vm^ or bifunctional degraders recruiting VHL E3 ligase. No signal was observed upon treatment with inactive control molecules (Figures 3B and C and S3c).

**Figure 3.**
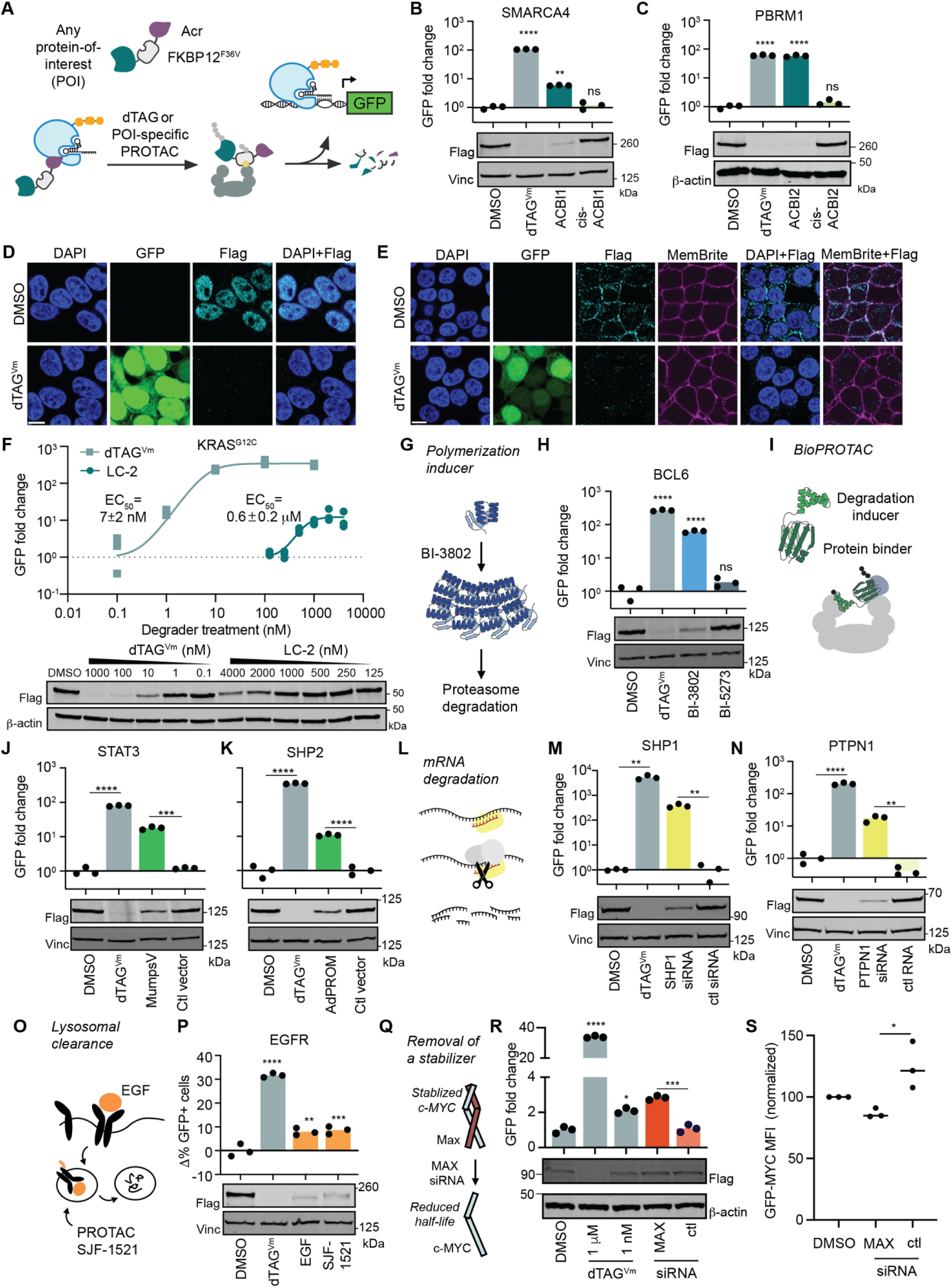
RTA circuits function with various target proteins and are agnostic to mechanism of protein downregulation. (**A**) A Flag tagged protein-of-interest (POI) is fused to a degradation control protein (FKBP12^F36V^, degraded by dTAG^Vm^) and Acr for transcriptional sensing of protein degradation. (**B**) Quantified live cell fluorescence microscopy and representative immunoblot of the monoclonal SMARCA4-sensing line treated for 24 hours with DMSO, or 1 μM dTAG^V^, ACBI1, or cis-ACBI1. (**C**) Quantified live cell fluorescence microscopy and representative immunoblot of the monoclonal PBRM1-sensing line treated for 24 hours with DMSO or 100 nM dTAG^V^, ACBI2, or cis-ACBI2. Representative immunofluorescent confocal microscopy for **(D)** SMARCA4-sensing cells and (**E)** KRAS^G12C^-sensing cells, treated for 24 hours with DMSO or 1 μM dTAG^V^. (**F**) Quantified live cell fluorescence microscopy and immunoblot of the monoclonal KRAS^G12C^-sensing line treated with DMSO or increasing concentrations of dTAG^Vm^ and LC-2. EC_50_ values and standard errors are calculated from a four-parameter logistic equation curve fit. (**G**) BI-3802 polymerizes BCL6 and promotes degradation. (**H**) Quantified live cell fluorescence microscopy and immunoblot of the monoclonal BCL6-sensing line treated for 24 hours with DMSO, 1 μM dTAG^Vm^, BI-3802, or BI-5273. (**I**) Synthetic (AdPROM) and viral (mumps virus V protein: mumpsV) bioPROTACs engage E3 ligases to degrade their targets. Quantified live cell fluorescence microscopy and immunoblot of a monoclonal (**J**) STAT3-sensing line treated for 24 hours with DMSO, 1 μM dTAG^Vm^, mumpsV transfection, or control transfection, and (**K**) SHP2-sensing line after 24 hours treatment with DMSO, 1 μM dTAG^Vm^, SHP2-AdPROM transfection, or control transfection, (n=3). (**L**) mRNA degradation by siRNA is a post-transcriptional protein downregulation mechanism. Quantified live cell fluorescence microscopy and immunoblot of the monoclonal SHP1-sensing line (**M**) and the PTPN1-sensing cell line (**N**) after 48 hours treatment with DMSO, 1 μM dTAG^Vm^, or siRNA transfection (n=3). (**O**) Lysosomal EGFR degradation is triggered by EGF ligand engagement as well as via direct C-terminal ubiquitination by PROTAC molecules. (**P**) Flow cytometry and immunoblot of a monoclonal EGFR-sensing line treated for 48 hours with DMSO, 1 μM dTAG^Vm^ or SJF-1521, or 500 ng/mL EGF protein. (**Q**) Inhibition or downregulation of biological stabilizers for target destabilization. (**R**) Quantified live cell fluorescence microscopy and immunoblot of a monoclonal MYC-sensing line treated for 48 hours with 1 μM dTAG^Vm^, 1 nM dTAG^Vm^, MAX siRNA, or control siRNA. (**S**) Flow cytometry of a GFP-MYC stable monoclonal cell line treated with MAX siRNA or control siRNA for 48 hours, normalized to DMSO. Statistics: Ordinary one-way ANOVA with Dunnett’s multiple comparison to DMSO (B, C, H, P, and R), and multiple unpaired two-tailed t tests (J, K, M, N, R (siRNA treatments), and S). Immunoblots in this figure detect Flag-tagged proteins-of interest and vinculin or β-actin loading controls, and live cell fluorescent microscopy data are depicted as GFP fold change normalized to DMSO controls (n=3).

SMARCA4-Acr localizes to the nucleus by confocal microscopy (Figure 3D), consistent with native SMARCA4 localization. KRAS^G12C^ contains a membrane-anchoring C-terminal farnesylation signal which dynamically engages with the plasma membrane or endosomal membranes.^32^ Acr-KRAS^G12C^ circuits exhibited high nonspecific GFP production that was mitigated when Cas was expressed from a weak (PGK) promoter, enabling the selection of monoclonals (Figure S3d-g). PGK-promoted dCasA parental cells with Acr-KRAS^G12C^ exhibited primarily non-nuclear KRAS^G12C^ by confocal microscopy (Figure 3E). Both dTAG^Vm^ and LC-2, a covalent KRAS^G12C^ PROTAC, led to degradation of KRAS and GFP production (Figure 3F).

RTA by degradation should be agnostic to the mechanism of protein-Acr fusion downregulation and the molecular identity of the degrader. To test this, we chose the BCL6-BI3802 target-degrader pair which operates by a unique, small molecule-induced polymerization-coupled degradation mechanism (Figure 3G and S3h)^33^. Like other degraders tested, we were able to observe signal activation and degradation with both dTAG^Vm^ and BI-3802, but not with BI-5273, a negative control analog (Figures 3H and S3i). We also observed transcriptional activation using both virally-derived (mumps virus V protein)^34^ and synthetic (SHP2-AdPROM)^35^ protein-based degraders of STAT3 and SHP2, respectively (Figures 3I-K and S3j). Finally, to demonstrate that target transcript downregulation can also lead to RTA, we built cell circuits containing SHP1-and PTPN1-Flag-FKBP12^F36V^-AcrIIA4 fusions and demonstrated that siRNA-mediated mRNA depletion results in decreased protein levels by immunoblot and substantial activation of GFP expression. (Figures 3L-N and S3k).

EGF treatment can induce ubiquitination, internalization, and subsequent lysosomal degradation of EGFR.^36^ EGFR-targeted PROTAC molecules, such as SJF-1521, lead to ubiquitination and EGFR shuttling through the same lysosomal degradation pathway (Figure 3O). To assess if we could detect protein degradation through the lysosome, we examined EGFR downregulation. We generated a polyclonal cell line utilizing high expressing EF1α-driven EGFR to reduce background GFP fluorescence, which showed poor signal induction (Figure 3Sl). This circuit was likely limited by the enforced separation between nuclear VPR-dCas9 and plasma membrane EGFR-Acr. Multiple rounds of FACS sorting with 100 nM dTAG^Vm^ treatments resulted in a monoclonal cell line that demonstrated an increase in reporter signal upon degradation by EGF and SJF-1521 by flow cytometry (Figure 3K and 3Sl), but a lack of sensitivity is potentially inherent in the forced separation of the components by localization, requiring large Acr/dCasA ratios (Figure S3m) and measurement by flow cytometry due to the small change in GFP production.

Ideally, RTA-based abundance sensing could also inform on pathway-induced destabilization of a target protein for biological study or therapeutic discovery. To test this, we transfected a monoclonal MYC RTA cell line (Figure S3n) with siRNA against its stabilizing heterodimer partner MAX (Figure 3Q).^37^ Upon knockdown of MAX we observed circuit activation, whereas the corresponding protein degradation effect was not clear by immunoblot as a function of near-native levels of MYC incorporation (Figure 3R). To confirm the observed effect was both on-target and subtle, we generated a GFP-MYC fusion and showed destabilization upon MAX siRNA treatment by flow cytometry (Figure 3 and S3o). The development of RTA circuits for nine disease-relevant proteins reveals general RTA principles: nuclear and many cytoplasmic target circuits respond best with strong dCasA expression, whereas proteins with more limited dCasA access show improved performance at higher Acr/dCas9A promoter strength ratios. Combined, these data demonstrate sensitive detection of both direct and network-mediated protein destabilization across diverse target proteins.

### Multiplexed detection of protein degradation

Multiple protein-to-transcription computations are performed simultaneously in cells to create complex regulatory and response systems. To generate two orthogonal synthetic RTA circuits that operate simultaneously in a single cell, five components must be integrated: guideRNAs, Cas proteins, Acrs, transcription promoters, and target genes (Figure 4A). Both *Staphylococcus aureus (Sau)* Cas9 and *Lachnospiraceae bacterium (Lb)* Cas12a have orthogonal PAMs and sgRNA structures to *Spy*Cas9.^38,39^ Additionally, both can be inhibited by characterized anti-CRISPR proteins; AcrIIA13b is a truncated anti-CRISPR from *Staphylococcus schleiferi* that inhibits DNA binding by *Sau*Cas9,^40^ and AcrVA4 derived from *Moraxella bovoculi* inhibits *Lb*Cas12a.^41^ We generated a modified version of a previously reported *Sau*dCas9 CRISPRa construct.^42^ After stable integration of AcrIIA13b-iRFP713-FKBP12^F36V^, GFP was produced upon dTAG^Vm^ treatment (Figure S4a). To avoid inactivation or loss of *Sau*dCas9, we engineered a construct with the antibiotic selection marker expressed on the same transcript separated by a self-terminating peptide (2A) and demonstrated that we could sense STAT3 degradation by flow cytometry (Figure 4B and S4b) and live cell fluorescence imaging (Figure 4C).

**Figure 4.**
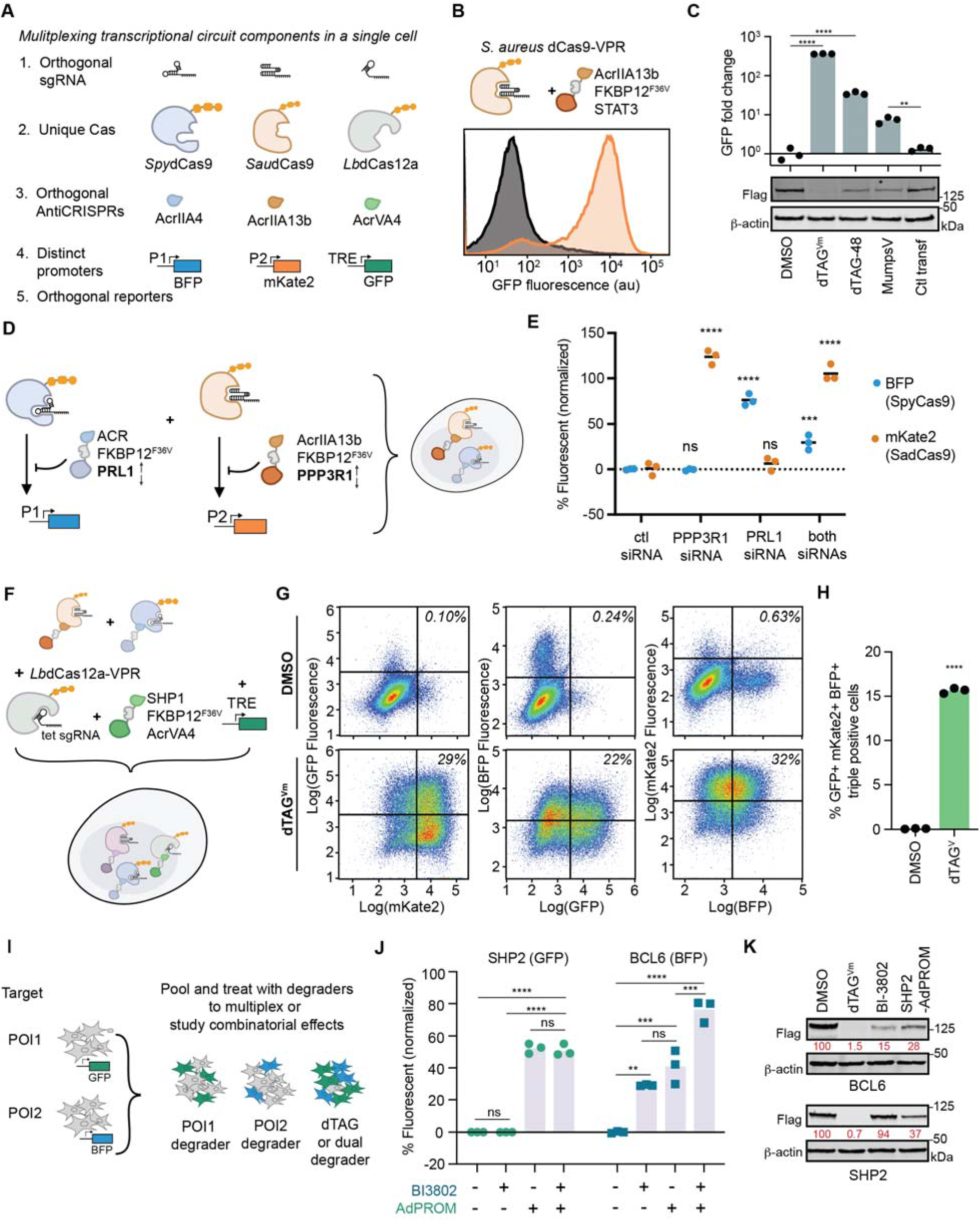
Multiplexing protein degradation sensing with multicellular pooled assays as well as orthogonal Cas/Acr pairs in a single cell. (**A**) Five circuit components must operate orthogonally to allow for multiple RTA circuits to function within a single cell. (**B**) Representative flow cytometry histograms of *Sau*dCas9 circuits detecting STAT3 protein levels upon treatment with DMSO or 1 μM dTAG^Vm^ for 48 hours. (**C**) Quantified live cell fluorescence microscopy and immunoblot of the monoclonal STAT3-sensing *Sau*dCas9 circuit cell line treated with DMSO, dTAG^V^, dTAG-48, or transfected with MumpsV or control vector after 48 h. (**D**) Schematic of dual orthogonal Cas circuit monoclonal cells stably expressing *Spy*dCas9 and *Sau*dCas9 with orthogonal sgRNAs, promoters, fluorophores, and target protein-Acr fusions (PRL1-AcrIIA4 and PPP3R1-AcrIIA13b). (**E**) Change in percentage of fluorescent cells was detected by flow cytometry (n=3) 48 hours after treatment with 1 μM dTAG^V^, or siRNAs transfection. (**F)** A triple orthogonal Cas/Acr monoclonal cell line created by addition of *Lb*dCas12a-VPR to the cells from (D) with an orthogonal sgRNA, GFP fluorescent readout, and protein-of-interest SHP1 fused to AcrVA4 anti-CRISPR. (**G)** Representative flow cytometry pseudocolor plots of dual fluorescent turn-on comparing DMSO to 1 μM dTAG^Vm^ after 72 hours. Total triple fluorescently positive cells are quantified in (**H**). (**I**) Degradation-sensing cell lines with orthogonal fluorescent outputs pooled into a single assay. (**J**) Triplicate flow cytometry of fluorescence activation upon treatment with individual and combined degraders in pooled wells of BCL6- and SHP2-sensing cell lines with orthogonal (BFP and GFP) outputs, normalized to max (1 μM dTAG^Vm^) and min (DMSO) for each cell line. (**K**) Representative immunoblot (n=3) of degradation of BCL6 and SHP2 by treatment with DMSO or 1 μM dTAG^Vm^ or BI-3802 for 24 hours, or transfection with SHP2-AdPROM for 48 hours. Statistics: One way ANOVA with Tukey’s multiple comparison test (B), or Dunnett’s multiple comparison to untreated/DMSO control (C, E), and unpaired two-tailed t test (C, comparing transfected samples, and J).

To test the ability to combine orthogonal circuits, we integrated the *Sau*Cas9 and *Spy*Cas9-based circuits into a single monoclonal cell line. Using two phosphatase subunits fused to orthogonal anti-CRISPR proteins as the targets (PRL1-AcrIIA4 and PPP3R1-AcrIIA13b, Figure 4D), we were able to demonstrate independent reporter production selectively upon knockdown of each target mRNA, as well as combined reporter activation upon treatment with dTAG^Vm^ or dual mRNA knockdown (Figures 4D and E and S4c). Finally, we integrated a *Lb*Cas12a^43^ circuit (Figure S4d) into these cells to generate a cell line which contained three independent ratiometric activation systems simultaneously (Figures 4F and S4e). We treated these cells with dTAG^Vm^ and observed simultaneous operation of all three circuits with orthogonal outputs (Figures 4G and H).

RTA also enables multiplexing in a pooled format via orthogonal outputs to sense degradation of numerous targets simultaneously. We isolated monoclonal cell lines that could detect degradation of SHP2 via GFP and BCL6 via mTagBFP (BFP). The cell lines were pooled and treated alone or in combination with their respective degraders (Figures 4I-K). BI-3802 specifically activated the BCL6 cells; whereas the bioPROTAC SHP2-AdPROM, designed to degrade SHP2 specifically, also led to BCL6 downregulation and circuit activation (Figures 4J and K and S4f). An additive effect of SHP2-depletion and BI-3802 treatment on BCL6 was detected by the assay (Figure 4J). SHP2-depletion has been reported to cause BCL6 downregulation by immunoblot in multiple lymphoma models,^44^ supporting our circuit-based observations.

### Modular outputs for cell survival and degradation detection by sequencing

Cells use protein degradation to trigger numerous phenotypically distinct outputs. In principle, RTA allows protein degradation to initiate production of any gene of interest. To demonstrate this, we built a circuit that upon degradation of target protein would result in production of a constitutively active caspase 3,^45^ leading to cell death (Figure 5A). We found this output predictably selected against leaky reporter expression in untreated cells, with control over cell death via target protein degradation at 48 h (Figure 5B). At later timepoints (∼72 hours) the fraction of cells within the polyclonal population that fail to activate cytotoxic response upon protein degradation dominate (Figures 5C and S5a). Additionally, we built a circuit that could transform the degradation of a target protein into the production of an antibiotic resistance gene (blasticidin S deaminase) (Figure 5D). Survival of cells after treatment with blasticidin would indicate target degradation. We observed an increase in cell confluency upon cotreatment with dTAG^Vm^ and blasticidin (10 μg/mL) at 48 hours (Figure 5E) and 72 hours (Figures 5F and S5b).

**Figure 5.**
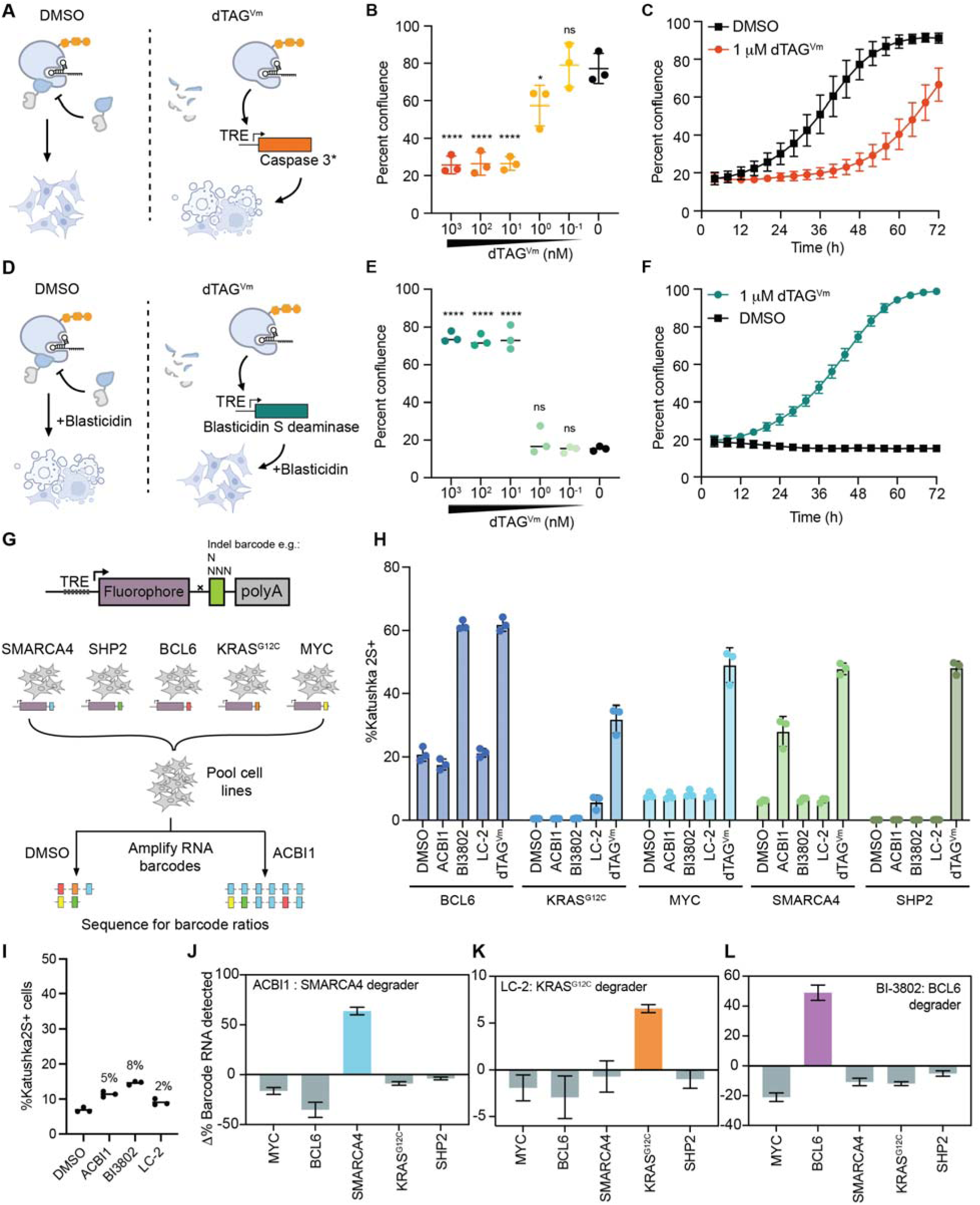
Protein degradation can drive modular reporter outputs. (**A**) Cell lines with a constitutively active mutant caspase3 reporter gene will induce death upon circuit activation. (**B**) Percent confluence detected by live cell microscopy of cells with a caspase reporter gene after 48 hour treatment with a range of dTAG^Vm^ concentrations. (**C**) Percent confluence over time was detected by live cell fluorescence microscopy for the DMSO and 1 μM dTAG^Vm^ treatments in (B). (**D**) Cell lines with the blasticidin resistance gene (bsr) as the reporter gene resist cell death when co-treated with blasticidin (10 μg/mL) and degrader molecule. (**E**) Percent confluence detected by live cell microscopy of cells with a bsr reporter after 48 hour treatment with a range of dTAG^Vm^ concentrations. (**F**) Percent confluence over time was detected by live cell fluorescence microscopy for the DMSO and 1 μM dTAG^Vm^ treatments in (E). (**G**) In addition to any reporter gene of interest (Katushka2S fluorophore) a 3’UTR RNA barcode can be included on the reporter gene. Cell lines containing barcoded RNA reporter genes can be multiplexed by pooling cell lines, treating with degraders, and sequencing can be used to deconvolute which proteins were downregulated. (**H**) Five polyclonal cell lines sensing degradation of SMARCA4, SHP2, BCL6, KRAS^G12C^, and cMYC were assessed for Katushka2S turn on individually by flow cytometry, 24 hours after treatment with DMSO or 1 μM treatment with dTAG^Vm^, ACBI1, BI3802, and LC-2 (n=3). (**I**) Flow cytometry from the pooled sequencing experiment described in (G) 24 hours after treatment with DMSO or 1 μM treatment with dTAG^Vm^, ACBI1, BI3802, and LC-2 (n=3). Changes in protein-of-interest-specific barcode sequences detected by sequencing upon treatment with (**J)** 1 μM ACBI1, (**K**) 1 μM LC-2, or (**L**) 1 μM BI-3802 compared to DMSO at t=24 hours (n=3).

The ability to transform protein-level information into nucleic acid readouts has immense potential for scalable, rapid, sensitive, and highly multiplexable study of post-transcriptional changes. We reasoned RTA could produce RNA barcodes that would facilitate multiplexed readout of protein degradation by sequencing (Figure 5G). To demonstrate this, we pooled five cell lines with distinct targets (SMARCA4, BCL6, SHP2, KRAS^G12C^, and cMYC) each containing a unique indel-based barcode incorporated polyclonally (Figures 5G and H and S5c). After pooling, the cell lines were treated with 1 μM ACBI1, BI-2802, LC-2, or DMSO control for 24 hours followed by flow cytometry (Figure 5I), cDNA generation and barcode amplification by PCR, and amplicon sequencing. We found that upon degrader treatment, the percentage of total reads for each distinct target-degrader combination increased dramatically compared to the DMSO-treated control pool (Figures 5J-L). Additionally, statistically meaningful degradation could be detected, and target deconvoluted using RNA barcode sequencing, exemplified by the robust signal for SMARCA4, KRAS^G12C^, and BCL6 degradation within the pool (Figures 5J-L). Thus, transforming target protein degradation into an amplified sequencing readouts increases both the multiplexability and sensitivity of RTA.

### Ratiometric transcriptional activation operates at endogenous protein levels

Since RTA decouples absolute target protein abundance from degree of signal activation, it may be uniquely useful in detecting low-expressing or endogenous protein fluctuations. To test this and to benchmark RTA against other live-cell protein detection mechanisms (direct fluorophore and HiBiT fusions), we used the PITCh knock-in protocol^46^ (Figure 6A) to insert RTA, fluorescent, and luminescent reporters fused to FKBP12^F36V^ at the N-terminus of five proteins chosen for diverse expression profiles^47^ and prior knock-in success^48^ in HEK293T cells (ATM, PRMT9, RIOK3, MEK1, and PARP1) (Figure 6B). After knock-in of the sensor protein, cells were engineered with reporter partner (dCasA, mNG1-10, or LgBit) and monoclonal cells were selected for all lines. Notably, engineering of RTA circuits requires no additional steps beyond those utilized for live cell HiBiT. Split neon green polyclonal knock-in cells were consistently dimmer than their GFP-fused counterparts (Figure S6a), thus only direct GFP fusions were selected for subsequent fluorophore-fusion comparisons. GFP fusion fluorescence intensities were consistent with the expected relative ordering of target protein abundance (Figure 6C). As this panel exhibited a range of Acr levels, we used EF1α-driven dCasA for the PARP1 cell lines, and sv40-driven dCasA for less abundant targets (Figure 6D). We observed that while Cas expression is higher in the PARP1 target line (Figure 6E), both PARP1 and ATM cell lines have Acr/Cas transcript ratios <1 (Figure 6F).

**Figure 6.**
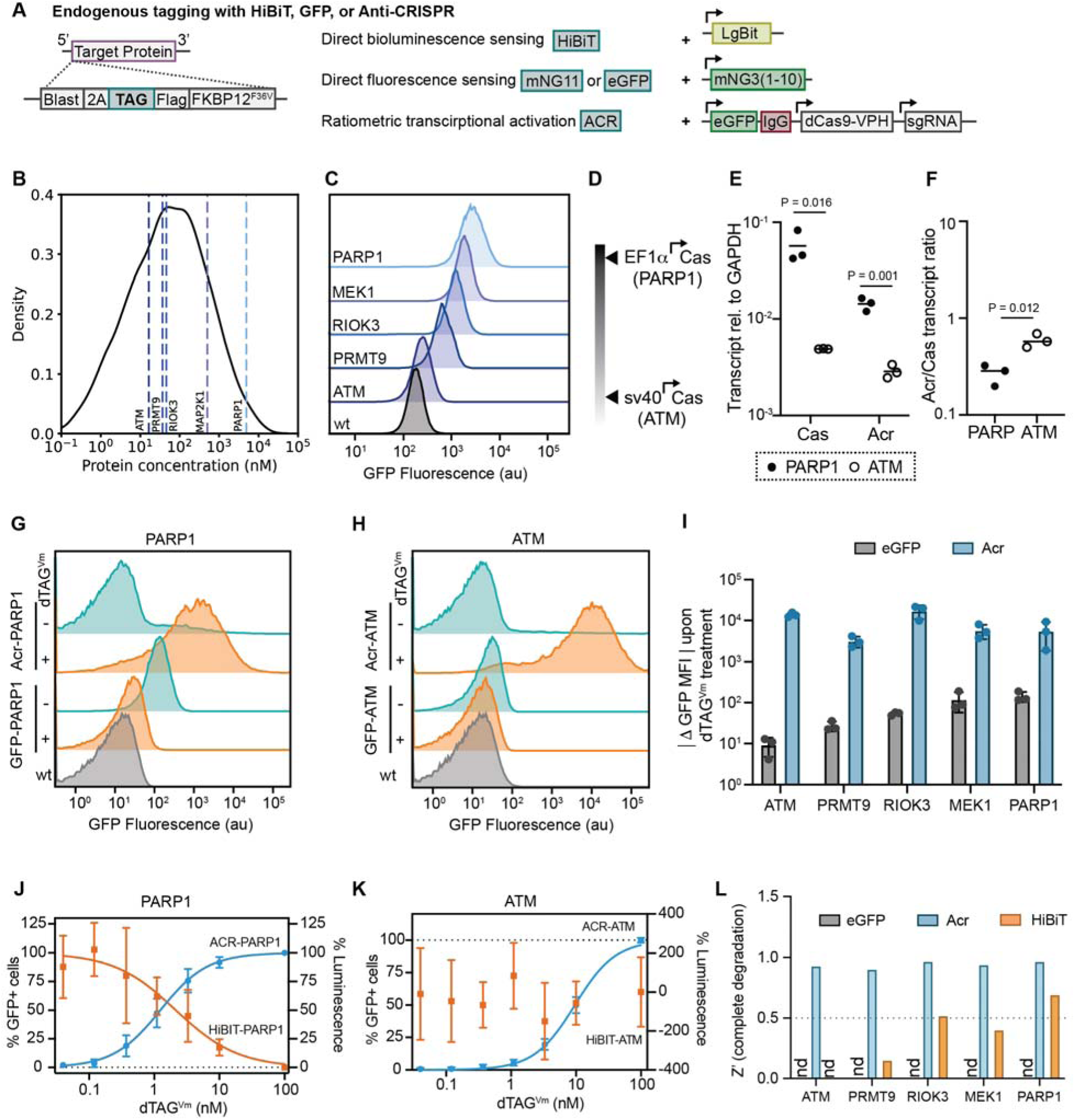
RTA circuits sense endogenous protein levels. (**A**) A PITCh knock-in protocol was used to N-terminally insert HiBiT, eGFP, mNG(11), and Acr into proteins, along with FKBP12^F36V^ as a positive control for inducible protein degradation. (**B**) Protein concentrations of the five knocked-in proteins, ATM, PRMT9, RIOK3, MEK, PARP1 reported by the OpenCell database. (**C**) Flow cytometry of eGFP knock-ins of the five endogenously tagged cell lines. (**D**) After knock-in of Acr to endogenous proteins, dCas9A was tested at different expression levels. A monoclonal containing EF1α-promoted Cas had the best sensitivity for PARP1, sv40-pormoted dCas9A were chosen for the other cell lines. (**E**) dCas9 and Acr transcript levels for the highest and lowest-expressing protein targets (PARP1 and ATM) were assessed by qPCR. (**F**) Acr/dCas9 ratios of PARP1 and ATM cell lines determined by qPCR. Flow cytometry of the cell line with the lowest concentration endogenous tagged protein (ATM) (**G**) and the highest (PARP1) (**H**) treated with DMSO or 100 nM dTAG^Vm^ for 24 hours. (**I**) Summarized flow cytometry of all eGFP and Acr RTA lines showing the change in GFP detected (ΔGFP) upon 100 nM dTAG^Vm^ treatment compared to DMSO. Magnitude of change is shown for comparison so eGFP change is positive. Live cell fluorescent confocal microscopy and plate reader luminescence for ATM (**J**) and PARP1 (**K**) treated with varying dTAG^Vm^ concentrations for 24 hours. (**L**) Z’-scores quantified for high-throughput 384-well format analysis of eGFP, Acr, and HiBiT tags for the endogenous proteins

Flow cytometry of GFP-fusion lines treated with dTAG^Vm^ showed that extent of signal changes upon target depletion were directly correlated with protein expression level (Figures 6G-I). However, as RTA utilizes transcription, extent of fluorescence changes upon degradation were decoupled from expression level of Acr-POI (Figures 6G-I and S6b). We sought to compare detection methods utilizing a 384-well plate format to measure changes in signal by high-throughput confocal microscopy for fluorophores and plate reader luminescence for HiBiT. However, direct GFP fusions to endogenous proteins were not differentiable from background by live cell, high-throughput confocal microscopy. Comparative analysis of dose response curves for degradation between RTA and HiBiT systems revealed that while both could detect near complete loss of PARP1 protein, only RTA was able to differentiate moderate amounts of degradation (Figure 6K). For ATM in the high-throughput live-cell context, HiBiT was unable to detect degradation while RTA exhibited similarly robust detection as that observed for PARP1 (Figure 6I). Across our panel of expression levels, only RTA exhibited Z’ scores greater than 0.5 for all proteins, indicating a highly robust live-cell high-throughput assay (Figure 6K and S6c).

### Ratiometric transcriptional activation for high-throughput small molecule profiling

Sensitive detection of protein degradation in pools and in high-throughput screening contexts could be particularly powerful for discovery of small molecule probes and therapeutic leads. One approach for targeting proteins that are canonically difficult to engage with small molecules is to target their proteostasis network or to target their degradation with molecular glues.

MYC is a master oncogenic regulator that has been indirectly targeted through a number of strategies.^49^ Macrolides such as FK506 destabilize MYC via calcineurin inhibition leading to altered phosphorylation status.^50^ We wondered whether high-throughput analysis of bioactive molecule libraries would enable direct observation of these destabilizing phenomena. Our MYC RTA cell line was well-suited for high-content imaging compared to a direct GFP-MYC fusion (Figures 7A, B and S7A-C), proving highly sensitive compared to triplicate immunoblot and maintaining robust reproducibility in a high-throughput screening context, even in combination with an orthogonal readout cell line (Figures 7C and D and S7d). We utilized a library consisting of 7,479 bioactive, clinically investigated molecules (Figures 7E), and screened RTA cells containing MYC, STAT3, FOXP3, and SHP2 as potential targets (Figures 7F and G). Of note, the FOXP3 line exhibited the highest sensitivity, rendering it a control group for non-specific circuit activation (Figure S7e). We found two features that enabled a high-true positive rate for these screens. The first was that simultaneous orthogonal viability and reporter measurements enabled clear differentiation between toxicity and destabilizers. The second was that the ability to multiplex targets in the same screen enabled rapid exclusion of non-specific effects, for example from global circuit activation by BET inhibitors (Figure S7f). Across this clinical drug library, we identified that the top three activating molecules for MYC destabilization were all Tacrolimus (FK506) accumulated from three different source libraries, and related macrolides also appeared as top hits (Pimecrolimus, Ascomycin). These molecules were validated independently for circuit activation (Figure S7g) and destabilization of a MYC-GFP fusion by flow cytometry with ∼20% loss detected (Figure 7H). Thus, RTA enables rapid hit identification of protein destabilizers from high-throughput screens.

**Figure 7.**
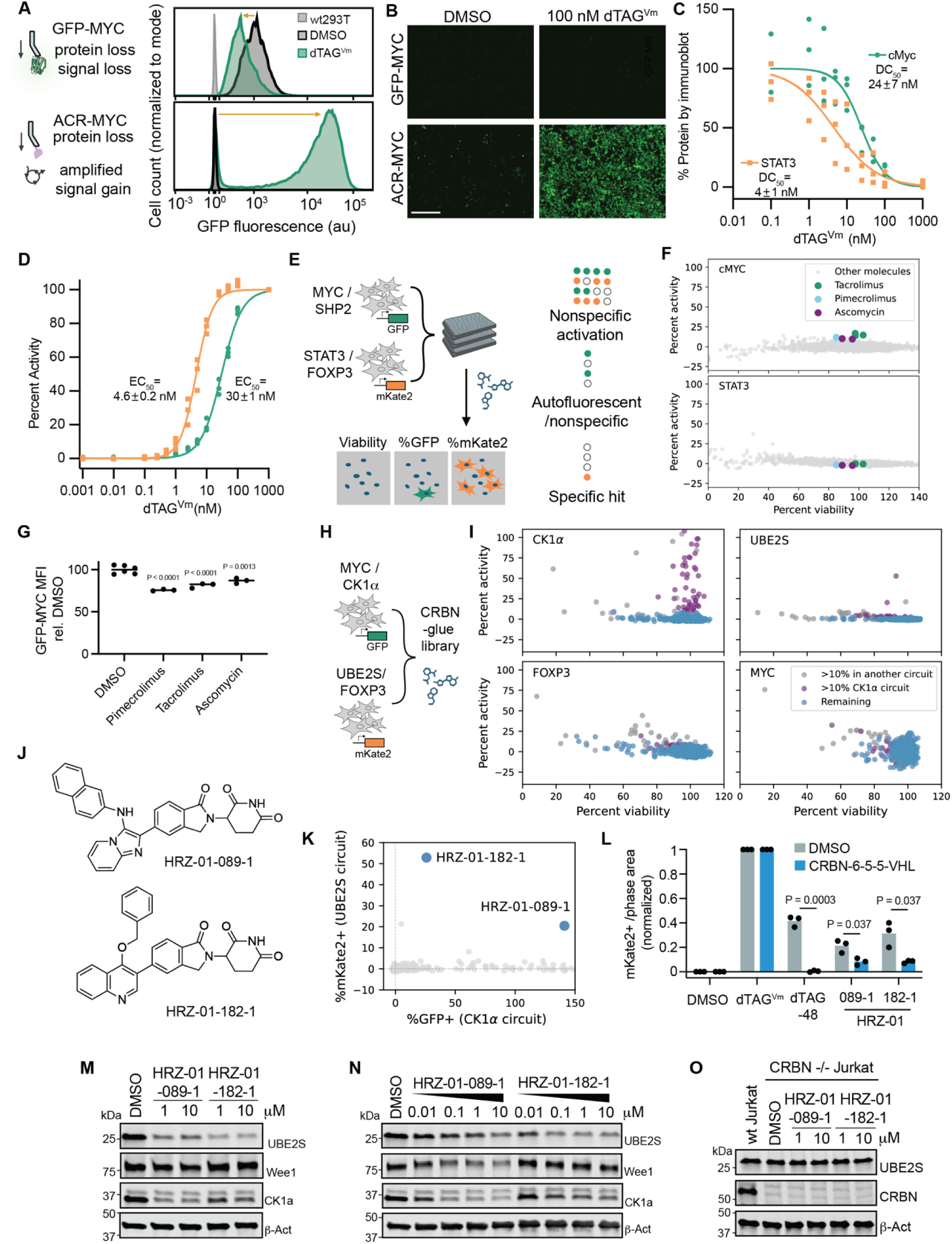
Multiplexing RTA circuits for high-throughput small molecule discovery. (**A**) Representative (n=3 flow cytometry histograms of GFP-MYC monoclonal cells (above) and Acr-MYC RTA circuits (below) upon DMSO or 100 nM dTAG^Vm^ treatment for 24 hours. (**B**) Live cell fluorescent microscopy GFP channel images comparing GFP-MYC fusion to Acr-MYC RTA circuits upon DMSO or 100 nM dTAG^Vm^ treatment for 24 hours. (**C**) Quantification and DC_50_ values from triplicate immunoblot of AcrIIA4-Flag-FKBP12^F36V^-cMYC and STAT3-Flag-FKBP12^F36V^-AcrIIA4 treated with various dTAG^Vm^ concentrations for 24 hours. (**D**) Percent fluorescent cell quantification by ImageXpress Micro Confocal live cell microscopy of pooled MYC and STAT3 cell lines in 384 well format after 24 hours treatment (n=3 replicates, ≥ 1 week apart, normalized to 1 μM dTAG^Vm^ and DMSO as max and min, respectively). (**E**) For high-throughput molecular profiling cells are seeded onto 384 well plates for 24 hours, compounds are added, 24 hours later, Hoechst dye is added and cells are visualized by ImageXpress Micro Confocal microscopy automated imaging. A clinical collection of 7,479 molecules was screened. Percent of mKate2+ and GFP+ cells compared to cell viability are reported. (**F**) MYC and STAT3 screen results of the clinical collection screen (n=1 replicate). (**G**) GFP-MYC MFI (normalized) to DMSO detected by flow cytometry after 24 h treatment with 10 μM tacrolimus, pimecrolimus, and Ascomycin. (**H)** A CRBN-glue containing library was screened against MYC, CK1α, FOXP3, and UBE2S, results are shown in (**I)** plotting percent activation of each circuit compared to cell viability in the well. Molecules that showed >10% activity in multiple circuits are in gray, molecules that showed >10% activity in CK1α are in purple. (**J)** Chemical structures of HRZ-01-089-1 and HRZ-01-182-1. (**K)** Plotting (n=1) screening results of the CRBN-glue library against UBE2S and CK1α (**L)** UBE2S RTA circuits were pretreated for 2 hours with CRBN degrader 6-5-5, followed by addition of 100 nM dTAG^Vm^, dTAG-48, or 10 μM HRZ compounds. Circuit activation was detected by live cell fluorescence microscopy after 24 hours and results are normalized to dTAG^Vm^ (1) and DMSO (0) for both DMSO and 6-5-5 treated cells. (**M**) K562 cells were treated with 1 and 10 μM of HRZ molecules for 16 hours, then UBE2S, CK1, Wee1, and β-actin levels were assessed by immunoblot. (**N**) Jurkat cells were treated with 0.01, 0.1, 1, and 10 μM of HRZ molecules for 16 hours, then UBE2S, CK1, Wee1, and β-actin levels were assessed by immunoblot. (**O**) CRBN -/- Jurkat cells were treated with HRZ molecules at 1 and 10 μM for 16 hours, followed by UBE2S, CRBN, and β-actin detection by immunoblot.

Molecular glue degraders enable depletion of target proteins without the need for high-affinity ligands. The revelation of consensus motifs recognized by certain E3 ligases, such as Cereblon (CRBN), has expanded the theoretical scope of targets which might be accessed via a unique glue degrader.^51,52^ Importantly, interactomic approaches have sought to find molecules which enforce E3-target engagement, but these approaches often do not detect degradation.^53^ Luminescence-based screening has been shown to result in a large number of false positives and remains challenging to multiplex, limiting the ability to identify true hits in a primary screen.^18^ We reasoned that RTA could enable rapid, multiplexed screening of glue-oriented libraries to discover new molecules for previously undegraded targets. We screened a custom library of 920 CRBN-targeting molecules against CK1α, a target previously shown to be degraded through CRBN hijacking.^54,55^ UBE2S, FOXP3, and MYC, which have not had direct molecular glue degraders disclosed (Figure 7H).^54^ The RTA screen resulted in many CK1α hits, as well as three hits for UBE2S (Figure 7I) an E2 protein known to enhance APC/C-mediated substrate priming.^56^ These UBE2S-active molecules originated from a combinatorial library previously developed for Wee1.^54^ Filtering these three for reduced activity in other RTA circuits resulted in two lead molecules with degradation activity against only UBE2S and CK1α (Figure 7I and J), with HRZ-01-182-1 appearing to be more potent and selective towards UBE2S (Figure 7K). We validated UBE2S circuit activation in independent RTA cell line experiments and demonstrated CRBN dependency through coincubation with the CRBN degrader (CRBN-6-5-5-VHL)^57^ (Figure 7L). Next, we confirmed that endogenous UBE2S was degraded by HRZ-01-089-1 and by the more potent HRZ-01-182-1 in both K562 (Figure 7M) and Jurkat (Figure 7N) cells. HRZ-01-182-1 also induced less degradation of CK1α and had negligible activity against WEE1 (Figures 7M and N). We further validated CRBN dependency through CRBN knockout in Jurkat cells and observed no degradation upon treatment with HRZ molecules (Figure 7O). The rapid identification of a new degrader against a protein not previously identified as being degradable establishes that RTA is a potent accelerator for discovery of degraders via high-throughput screening.

## Discussion

The global and selective turnover of proteins in cells defines circuits for modulating cellular functions and cycles. Modalities which seek to rewire these capabilities have the potential to disrupt these circuits in therapeutically-enabling ways. Sensing protein degradation in live cells enables an understanding of proteostasis as well as detection of therapeutic agents which modulate proteins and their proteostasis networks. We have developed a platform which converts the degradation of a protein of interest into any transcriptional output. This enables highly sensitive detection of protein degradation at scale, as well as programming of cellular function upon acute loss of a target protein. We develop the principle of ratiometric transcriptional activation by utilizing stoichiometric inhibitors (anti-CRISPR protein fusions) of targeted transcription factors (CRISPRa machinery). Perturbations in the levels of the Acr-protein fusion lead to active transcription of any gene of interest, with sensitivity defined by the ratio of Acr to Cas.

Programming genetic circuits to report on completely endogenous behavior with native component levels remains a challenge. Almost all synthetic reporters of protein degradation scale with expression level of target. Ratiometric transcriptional activation (RTA) enables sensing of degradation of low expression targets at endogenous levels via highly amplified readouts. We show the ability to sense many degradation mechanisms, suggesting broad applicability of RTA principles. Cells integrate multiple inputs to generate simultaneous but diverse outputs. We mimic this behavior with RTA by integrating multiple orthogonal circuits which sense changes in distinct proteins in a single cell. Practically, pooling individual cell lines with orthogonal reporters offers a way to rapidly multiplex. A challenge with pooling cells is both dilution of cells within the population as well the availability of unique and orthogonal outputs. We solve this conceptual challenge by directly linking the production of a unique nucleic acid barcode to the degradation of a target. This creates nearly limitless orthogonal reporters while providing an amplification mechanism for low-abundance reporters.

The identification and characterization of molecules that degrade targets has instigated pre-clinical programs directed against traditionally challenging drug targets. Few small molecules that degrade target proteins have been discovered prospectively using designer assays in high-throughput. We show that RTA, with the ability to multiplex, sense finer degrees of change, and amplify signal, provides a robust and uniform platform to discover target degrader or destabilizing molecules directly from screens. Detection of difficult or previously impossible to observe changes in protein levels directly in live cells at scale might reveal fundamentally new features of protein proteostasis networks.

### Limitations of the Study

RTA requires direct fusion of Acr to proteins-of-interest, which may alter their native function, localization, or interactions. For transmembrane proteins, the strong exclusion of the Acr from the nucleus can result in low sensitivity to degradation. Extensive single-cell sorting was needed to identify clonal cells that could operate in these contexts. We developed an all-in-one construct design to minimize engineering steps which proved highly sensitive even when deployed in polyclonal populations, however for certain applications single cell sorting or multiple integration events may be desired to capture a broader Acr/dCas9A range. We noticed that more sensitive RTA lines showed nonspecific activation upon BET protein inhibition, leading to potential false positives in small molecule screens. Multiplexing in these contexts was essential for identification of true positives through embedded counter screening. Due to transcriptional delay, RTA is not ideal for real-time protein level monitoring though it does provide a recording of a transient degradation event. Finally, we benchmarked RTA against other live-cell detection assays in a high-throughput 384-well plate screening context. As scale of experiment changes, alternative assays are likely to perform differently, though we note RTA retains statistical power at all scales attempted. We did not test alternative non-live cell approaches which may have improved sensitivities, primarily due to the advantage of directly measuring toxicity in screening contexts and biological experiments.

## Supporting information

Supplementary Figures, Methods, and Modeling

## Acknowledgments

Cell sorting/flow cytometry analysis for this project was done on instruments in the Stanford Shared FACS Facility. Certain data were collected on an instrument in the Shared FACS Facility using NIH Shared Instrument Grant 1S10OD026831-01. High-throughput chemical screening for this project utilized instruments from the Stanford High-Throughput Screening Knowledge center.

## Funding

Stanford Innovative Medicine Accelerator (DECS, MAG, SMB)

Stanford Cancer Center (DECS)

Molecular Devices ImageXpress Confocal (SIG S10OD066899)

Gordon and Betty Moore Foundation (MAG, SMB)

Stanford’s Knight-Hennessy Scholars Program (QQO)

Stanford Medical Scientist Training Program (NIH T32GM007365) (QQO)

Burroughs Wellcome Fund Careers at the Scientific Interface Award (MAG, SMB)

ChEM-H Chemistry-Biology Interface Training Program (JAC, KLR, MTT, QQO)

Stanford Medical Scholars Research Fellowship Program (QQO)

Stanford Undergraduate Summer Research Fellowship in Chemistry (AZX)

EDGE: Enhancing Diversity in Graduate Education Doctoral Fellowship Program (JAC)

Damon Runyon Cancer Research Foundation (SMC)

NSF Graduate Research Fellowships program (SAR)

Stanford CMAD Fellowship (SAR)

## Author contributions

Conceptualization: MAG, SMB

Methodology: MAG, JAC, KLR, MTT, AZX, SWC, HR, DESC, SAR, CAS, JY, SMB

Investigation: MAG, JAC, KLR, MTT, AZX, QQO

Visualization: MAG, SAR, JAC

Formal analysis: MAG, DESC

Funding acquisition: SMB

Supervision: NSG, SMC, SMB

Writing – original draft: MAG, SMB

## Data and materials availability

All data are available in the main text or the supplementary materials. Sequences of all plasmid inserts are provided in the supplementary information.

## Supplementary Materials

Materials and Methods

Figures S1 – S7

Supplementary References (*1-7*)

