## Supplementary Figures, Methods, and Modeling for "Ratiometric transcriptional activation by protein degradation"

**The PDF file includes:**

Figs. S1 to S7
Materials and Methods
Supplementary References

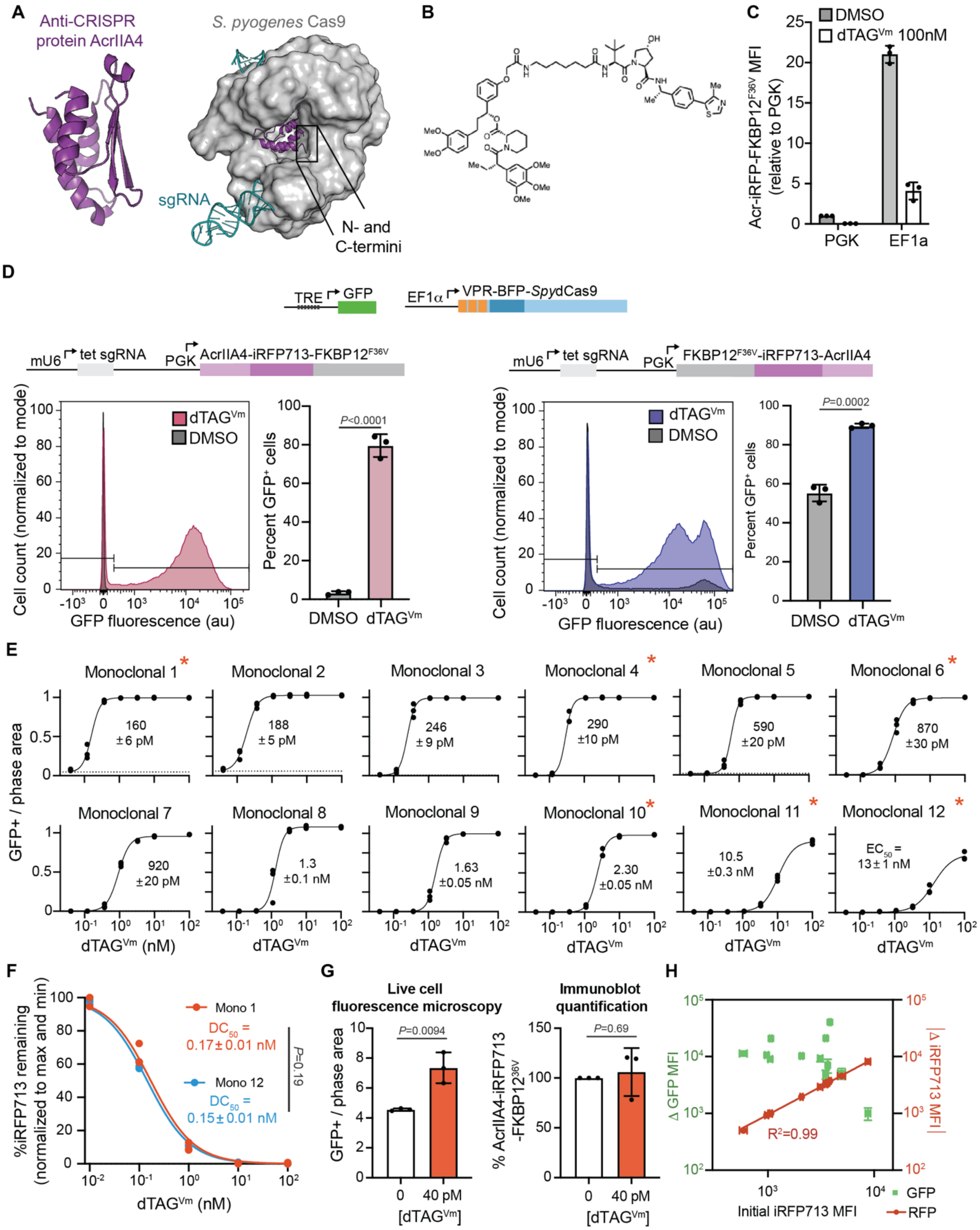


**Figure S1. A.** Structure of anti-CRISPR AcrIIA4 protein bound to SpyCas9 indicates that both the N- and C-termini are accessible for linkage to fusion proteins. (Left) AcrIIA4 anti-CRISPR protein from Listeria monocytogenes (PDB: 5XN4).[1] (Right) Crystal structure of AcrIIA4 bound to *Spy*Cas9 (PDB: 5VZL).[2] **B.**

Structure of the dTAG$^{V}$ analog dTAG$^{Vm}$ synthesized and used throughout this manuscript. dTAG$^{Vm}$ recruits the E3 ligase substrate adaptor VHL to induce ubiquitination and degradation to FKBP12$^{F36V}$ target protein. **C.** iRFP713 measurement by flow cytometry of the EF1$\alpha$- and PGK-promoted AcrIIA4-iRFP713-Flag-FKBP12$^{F36V}$ upon treatment with 100 nM dTAG$^{Vm}$ or DMSO control for 24 hours (mean$\pm$SD). **D.** Stable polyclonal cells with FKBP12$^{F36V}$ fused either N-terminally or C-terminally to Acr trigger GFP production upon dTAG$^{Vm}$ treatment. Polyclonal cells containing either A. FKBP12$^{F36V}$-Flag-iRFP713 fused to the N-terminus of AcrIIA4, or B. iRFP713-Flag- FKBP12$^{F36V}$ fused to the C-terminus of AcrIIA4, were treated with DMSO or 1 µM dTAG$^{Vm}$ for 24 hours and GFP fluorescence was measured by flow cytometry. Percent GFP+ cells and GFP MFI (mean$\pm$SD) are from n=3 independently performed experiments, with a representative histogram shown on the left. Statistics: unpaired two-tailed t tests. **E.** Percentage of GFP+ cells from twelve stable monoclonal cell lines (sorted in Figure 1G) are shown here after 24 hours treatment with different dTAG$^{Vm}$ concentrations by live cell fluorescence microscopy. $EC_{50}$ values and associated standard errors are calculated from a four-parameter logistic equation. Monoclonal curves noted with an asterisk* are also shown in Figure 1G. **F.** iRFP713 fluorescent protein loss by flow cytometry of monoclonal 1 and monoclonal 12 fit to a four-parameter logistic equation constrained to a maximum of 100 and minimum of 0 to calculate $DC_{50}$ and associated standard error of dTAG$^{Vm}$ treatment (normalized to max and min for each cell line). An unpaired two-tailed t test was used to compare the $DC_{50}$ values. **G.** The clonal cell line with the smallest $EC_{50}$ (Monoclonal 1) detects degradation of AcrIIA4-iRFP713-Flag-FKBP12$^{F36V}$ at 40 pM dTAG$^{Vm}$ (left) whereas protein degradation by triplicate immunoblot is not significantly detected (right). For the immunoblots, DMSO-treated wells were normalized to equal 100% for each replicate. Statistics: two-tailed unpaired t tests. **H.** Magnitude of the fluorescence change of iRFP713 loss (red circles with linear regression model fit) and GFP increase (green squares) by transcriptional activation for 12 monoclonal cell lines upon 1 µM dTAG$^{Vm}$ treatment by flow cytometry (mean ± SD, n=3). The x-axis depicts the relative target protein expression levels measured by iRFP713 fluorescence of DMSO-treated cells.

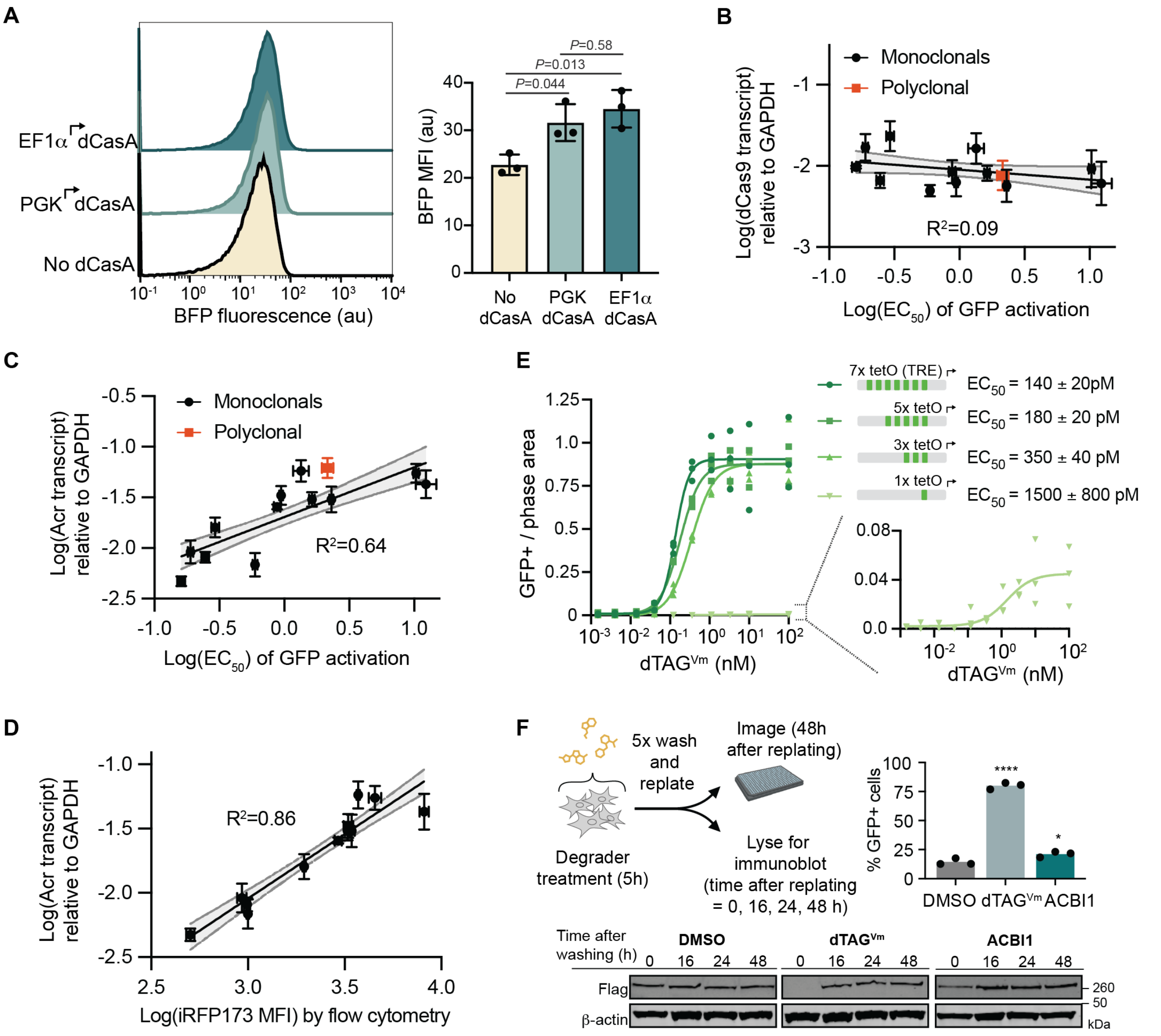


**Figure S2. A.** The VPR-BFP-*Spy*dCas9 construct stably integrated into cells via the Sleeping Beauty transposase system had minimal BFP fluorescence detected by flow cytometry under EF1α or PGK promoters. Data are from n=3 independently performed flow cytometry experiments (representative histogram on the left) and are analyzed with an ordinary one-way ANOVA followed by Tukey's multiple comparisons test. Transcript levels of (**B**) dCasA and (**C**) Acr in the 12 monoclonal cell lines (from Figure S1E) and the parent polyclonal line relative to GAPDH transcript were assessed by triplicate qPCR mean±SD. Linear fit with $R^2$ reported. **D.** Relative transcript levels of Acr correlated with a linear model to relative protein expression levels by iRFP713 detection via flow cytometry. **E.** tetO sites (dCasA sgRNA binding sites) at the promoter were varied from 1-7x and stably integrated as single RTA constructs into cells, GFP+ image area normalized to cell phase area of a polyclonal population was quantified by live cell fluorescent microscopy after 24 hours treatment with various $dTAG^{Vm}$ concentrations. The 1x tetO promoter is also shown as a magnified inset from 0% to 8% GFP+ activation on the y axis. **F.** SMARCA4-$FKBP12^{F36V}$-Acr RTA circuit cells were treated with 100 nM $dTAG^{Vm}$ or ACBI1 for 5 hours. Cells were then washed 5x with PBS, lifted with trypsin, and replated onto new plates for live cell fluorescent microscopy over time as well as timepoint immunoblot collection. GFP production at 48 hours after replating are shown, along with immunoblot analysis detecting Flag-tagged Acr construct and β-actin loading control over 48 hours.

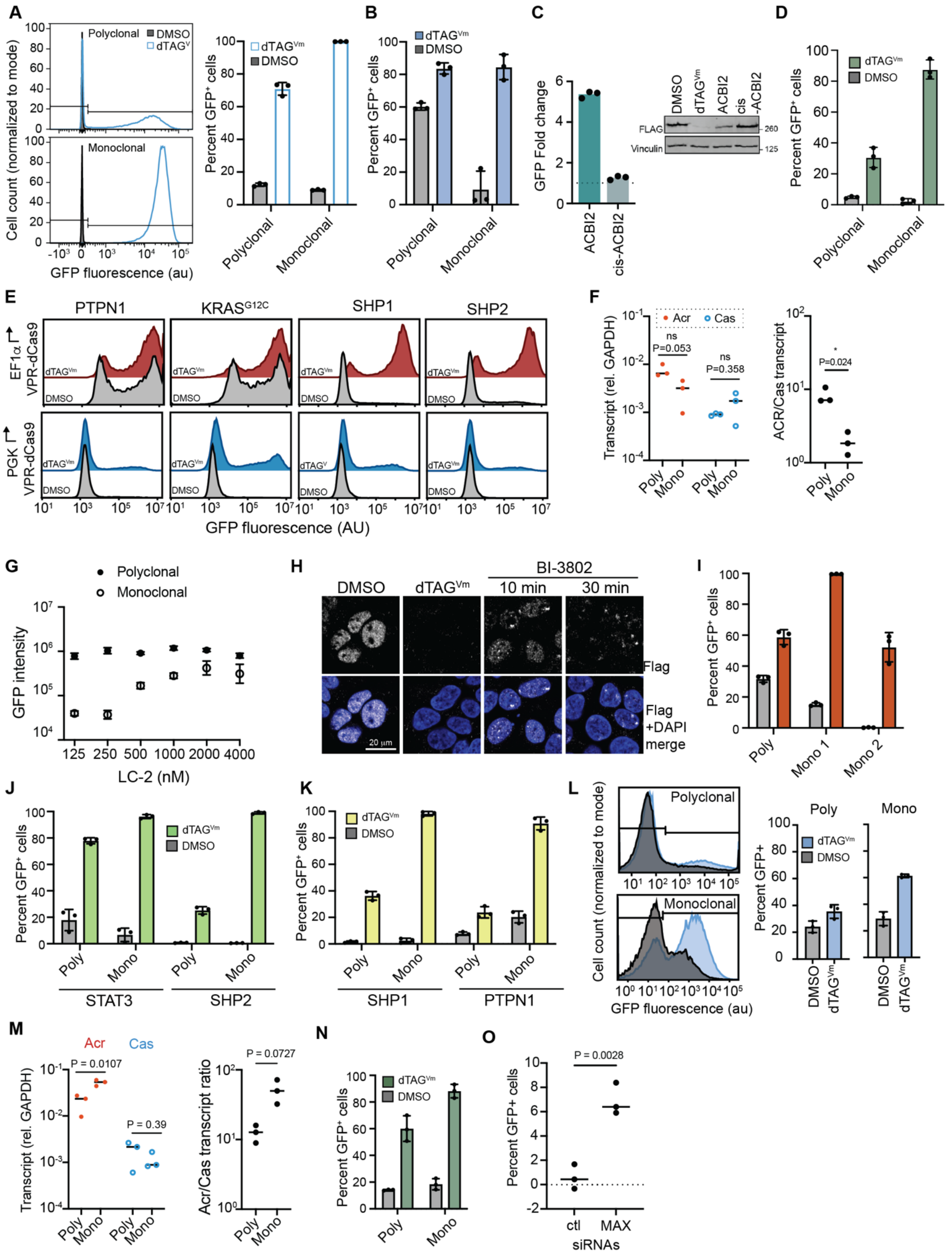

A
B
C
D
E
F
G
H
I
J
K
L
M
N
O
Polyclonal
Monoclonal
Cell count (normalized to mode)
GFP fluorescence (au)
Percent GFP+ cells
GFP Fold change
ACBI2
cis-ACBI2
DMSO
FLAG
Vinculin
PTPN1
SHP1
SHP2
GFP fluorescence (AU)
Transcript (rel. GAPDH)
Acr
Cas
ACR/Cas transcript
Poly
Mono
GFP intensity
LC-2 (nM)
BI-3802
10 min
30 min
Flag
Flag +DAPI merge
20 µm
Mono 1
Mono 2
STAT3
Acr/Cas transcript ratio
ctl
MAX
siRNAs

**Figure S3. A.** Polyclonal and monoclonal cell circuits detecting SMARCA4 degradation were treated with DMSO or 1 µM $dTAG^{Vm}$ and analyzed by n=3 flow cytometry experiments at t=24 h. Representative histograms on the left depict the gating used for the %GFP+ values quantified on the right. **B.** Polyclonal and monoclonal cell circuits detecting PBRM1 degradation were treated with DMSO or 1 µM $dTAG^{Vm}$ and analyzed by n=3 flow cytometry experiments at t=24 h. **C.** ACBI2 treatment of the monoclonal SMARCA4 circuit (previously characterized in Figure S3a) by live cell fluorescence microscopy, GFP integrated intensity was normalized to the DMSO average. Immunoblot (right) of SMARCA4 protein levels after 24 h treatment with either DMSO or 1 µM $dTAG^{V}$, ACBI2, or cis-ACBI2 control. **D.** Polyclonal and monoclonal cell circuits detecting KRASG12C degradation were treated with DMSO or 1 µM $dTAG^{Vm}$ and analyzed by n=3 flow cytometry experiments at t=24 h. **E.** Histograms for RTA circuits with either cytoplasmic target proteins (SHP1 and SHP2), or cytoplasmic-facing proteins that are anchored to membranes (PTPN1 largely ER-anchored) and ($KRAS^{G12C}$ largely PM-anchored) when treated with $dTAG^{Vm}$ degrader or control. Transcriptional background activity was lowered when circuits with membrane bound targets were developed with dCasA expressed on a weaker PGK promoter. **F.** Transcript levels of dCasA and Acr, as well as overall Acr/dCasA transcript ratio as assessed by qPCR are shown for the sensitive monoclonal $KRAS^{G12C}$ RTA circuit line compared to its parent polyclonal cells. **G.** $KRAS^{G12C}$ monoclonal and polyclonal GFP integrated intensity after 24 hour treatment with the LC-2 PROTAC at various doses and measured by live cell fluorescence microscopy. **H**. Confocal microscopy of Flag-tagged BCL6- RTA cells with a 1 µM BI-3802 polymerization-inducing degrader treatment for 10 min and 30 min. DMSO and 1 µM $dTAG^{Vm}$ treatment for 24 hours are shown as a negative and positive control, respectively. **I.** Polyclonal and monoclonal cell circuits detecting BCL6 degradation were treated with DMSO or 1 µM $dTAG^{Vm}$ and analyzed by n=3 flow cytometry experiments at t=24 h. Monoclonals can be selected for high ΔGFP% change (monoclonal 1) or optimized for high fold change (monoclonal 2). Polyclonal and monoclonal cell circuits detecting STAT3 and SHP2 degradation (**J**) and SHP1 and PTPN1 degradation (**K**) were treated with DMSO or 1 µM $dTAG^{Vm}$ and analyzed by n=3 flow cytometry experiments at t=24 h. **L.** Polyclonal and monoclonal cell circuits detecting EGFR degradation were treated with DMSO or 1 µM $dTAG^{Vm}$ and analyzed by n=3 flow cytometry experiments at t=48 h. Representative histograms on the left show the gating used for the %GFP+ values quantified on the right. The polyclonal and monoclonal lines for EGFR were analyzed at different days under different flow cytometer voltages and therefore gating is unique for each sample. **M.** Transcript levels of dCasA and Acr, as well as overall Acr/dCasA transcript ratio as assessed by qPCR are shown for the monoclonal EGFR RTA circuit line compared to its parent polyclonal cells. **N.** Polyclonal and monoclonal cell circuits detecting MYC degradation were treated with DMSO or 1 µM $dTAG^{Vm}$ and analyzed by n=3 flow cytometry experiments at t=24 h. **O.** MAX siRNA and control siRNA transfections into the MYC RTA circuits were assessed by flow cytometry 48 hours after transfection to confirm circuit activation by an independent method (normalized to 100 nM $dTAG^{Vm}$ and DMSO (max and min), respectively.

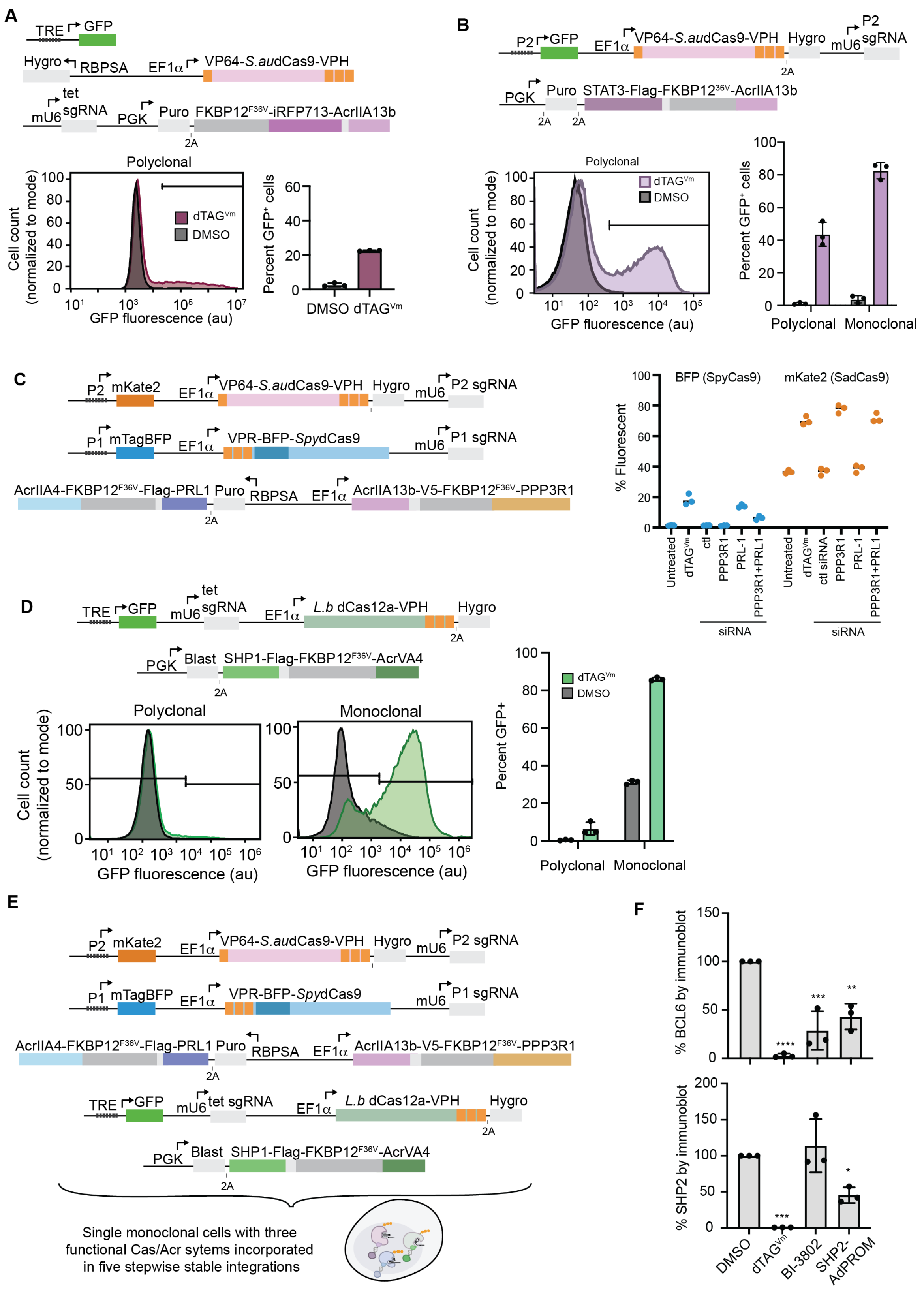


**Figure S4. A**. Composition of the three stably incorporated genes used to make the initial *S. au* dCas9 ratiometric transcriptional activation circuits in HEK293T cells. Polyclonal cell circuits detecting FKBP12$^{F36V}$

degradation were treated with DMSO or 1 µM dTAG$^{Vm}$ and analyzed by flow cytometry experiments after 24 hours of treatment (histogram shows n >10,000 cells). **B.** Characterization of polyclonal and monoclonal cell circuits utilizing VP64-*Sau*dCas9-VPH and its associated anti-CRISPR protein AcrIIA13b, fused to STAT3. Polyclonal *Sau*dCas9 cell circuits detecting STAT3 degradation were treated with DMSO or 1 µM dTAG$^{Vm}$ and analyzed by n=3 flow cytometry experiments at t=48 h. Representative histogram on the left show the gating used for the %GFP+ values quantified on the right. A representative histogram for the selected monoclonal circuit is shown in Figure 4B. **C**. Characterization of monoclonal cells containing dual orthogonal Cas RTA circuits. Three sequential exogenous gene insertions were used to make circuits that contained *Spy*dCas9/AcrIIA4 and *Sau*dCas9/AcrIIA13b ratiometric activation of transcription upon PRL1 and PPP3R1 phosphatase degradation, turning on BFP and mKate2, respectively. Raw data of triplicate flow cytometry experiments performed on independent days (normalized results shown in Figure 4E). **D.** Polyclonal and monoclonal *Lb*dCas9 cell circuits were treated with DMSO or 1 µM dTAG$^{Vm}$ and analyzed by n=3 flow cytometry experiments at t=48 h. Representative histogram on the left shows the gating used for the %GFP+ values quantified on the right. **E**. Five exogenous gene cassettes were sequentially incorporated into HEK293T cells to make three simultaneous orthogonal CRISPRa/Acr circuits detecting downregulation of each anti-CRISPR in a single cell. **F**. Quantification of triplicate immunoblot of two separate cell lines expressing BCL6-Flag-FKBP12$^{F36V}$-Acr and Acr-Flag-FKBP12$^{F36V}$-SHP2 degradation constructs each treated with DMSO, 1 µM dTAG$^{Vm}$, or 1 µM BI-3802 for 24 hours or transfected with SHP2-AdPROM for 48 hours. The first replicates are also shown in Figure 4C. Statistics: ordinary one-way ANOVA with Dunnett's multiple comparison test to DMSO.

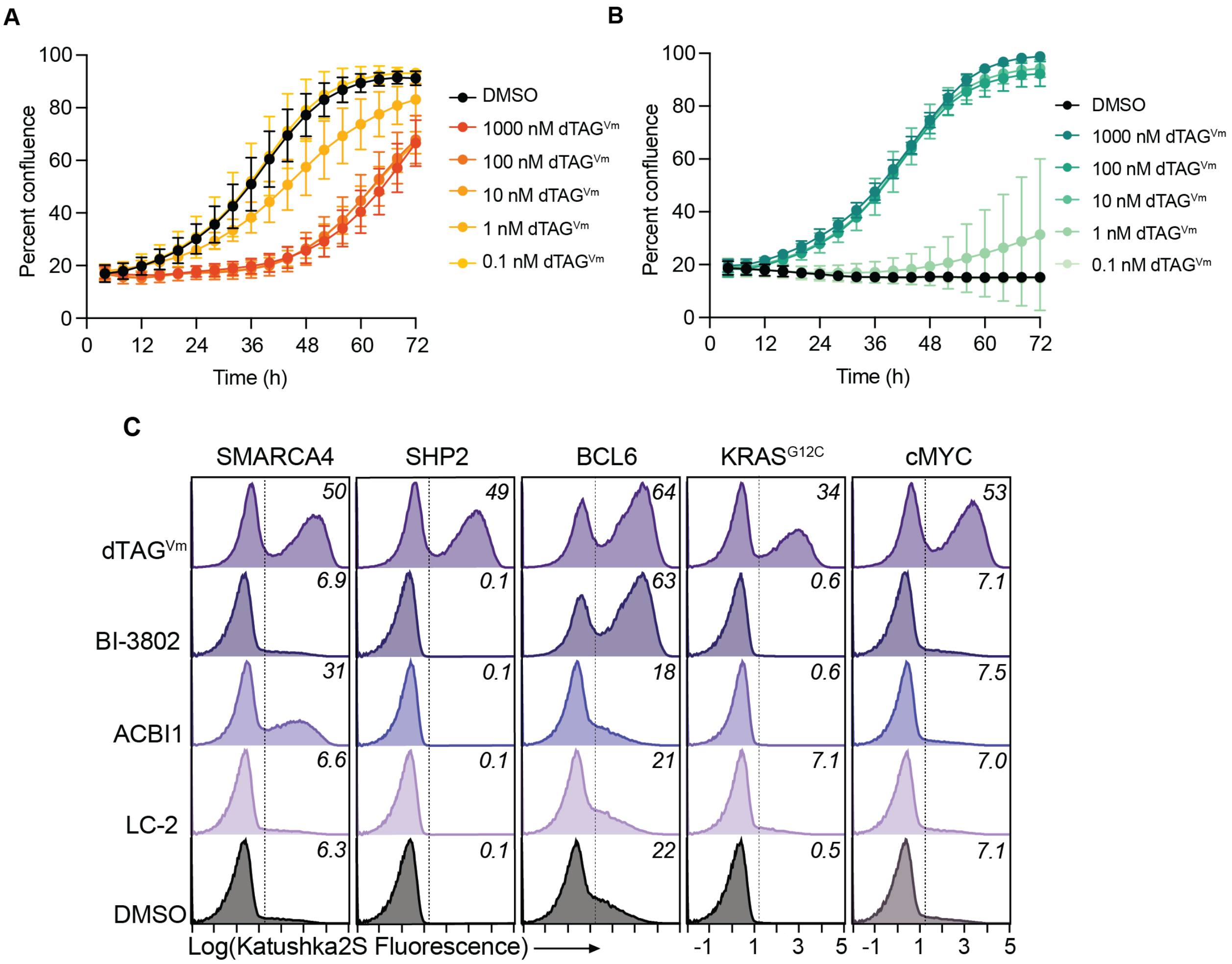


**Figure S5. A.** Percent confluence over time was detected for RTA circuits that induce death upon circuit activation by live cell fluorescence microscopy for DMSO and 1 µM – 0.1 nM dTAG$^{Vm}$ for three replicates performed on independent days (mean ± SD). **B.** Percent confluence over time was determined for RTA circuits with the blasticidin resistance gene (bsr) as the reporter by live cell fluorescence microscopy for co-treatments of blasticidin S (10 µg/mL) and either DMSO or 1 µM – 0.1 nM dTAG$^{Vm}$ treatments over 72 hours. Mean ± SD of 3 independently performed replicates are shown. **C.** Representative (n=3) histograms of five individual polyclonal cell lines sensing degradation of BCL6, KRAS$^{G12C}$, MYC, SMARCA4, and SHP2 assessed for Katushka2S turn on by flow cytometry. Gating for Figure 5H are shown as dashed lines.

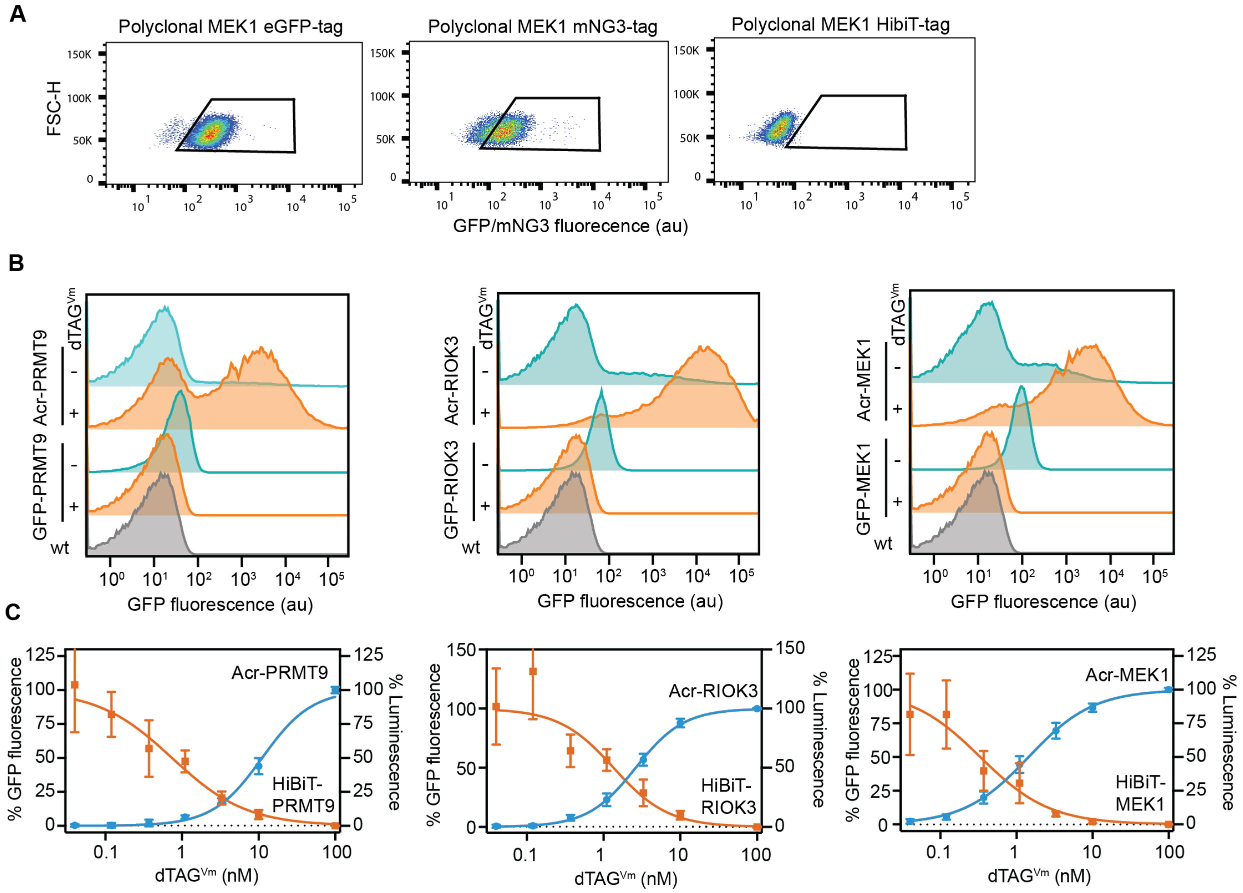


**Figure S6. A.** Flow cytometry showing gating for sorting monoclonal GFP+ cells. eGFP and mNG3 tagged cells were sorted for the GFP+ gate shown. HiBiT cells were sorted for single cells at random. **B.** Representative (n=3) flow cytometry experiments on PRMT9, RIOK3, and MEK1 cells comparing eGFP-tagged targets to ACR-tagged targets when treated with 100 nM dTAG$^{Vm}$. Quantified triplicate fluorescence changes are shown in Figure 6I. **C.** Triplicate 384 well plate analysis by luminescence (HiBiT) or high-throughput confocal microscopy (Acr) for PRMT9, RIOK3, and MEK1 tagged cells.

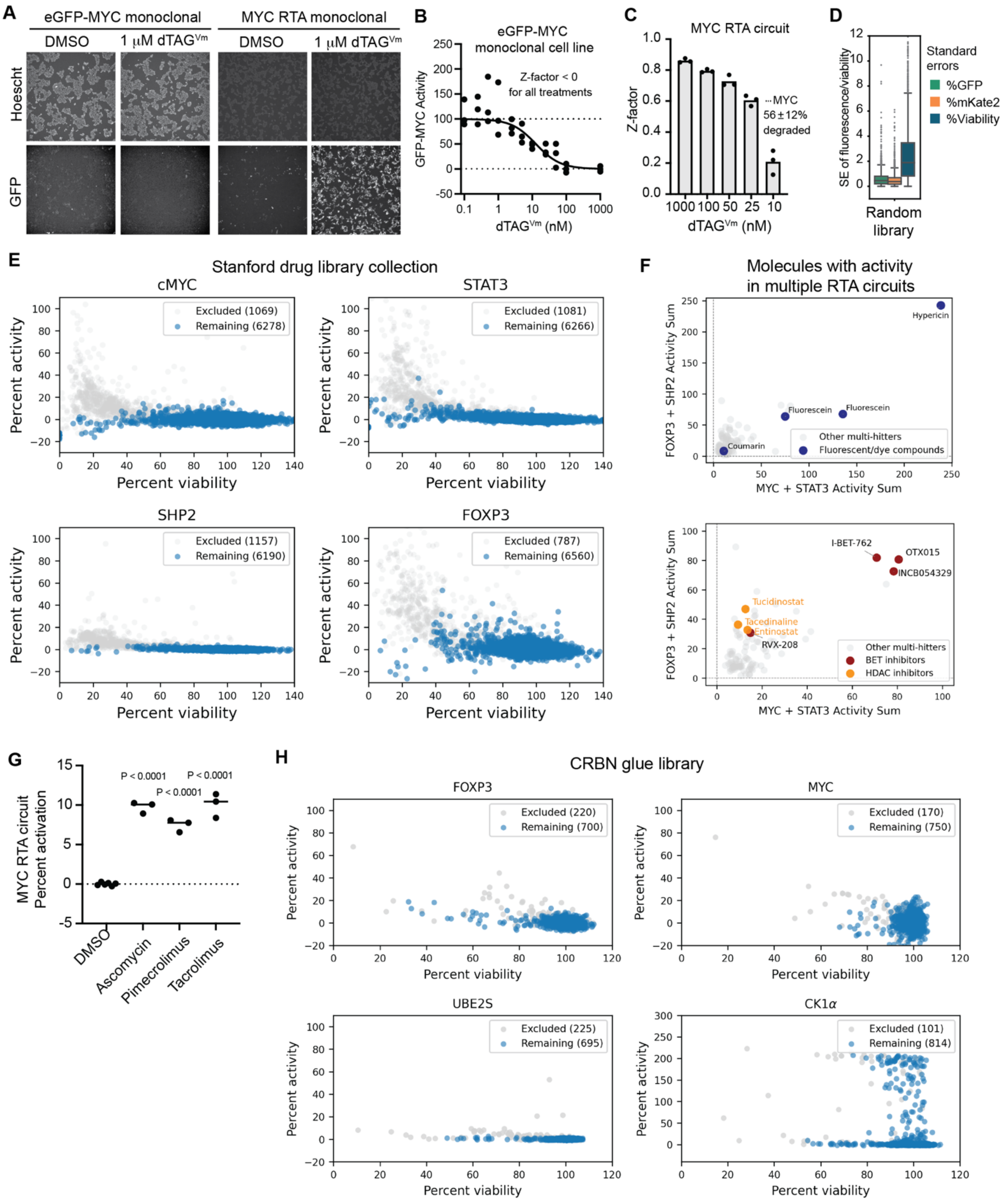


**Figure S7. A.** ImageXpress confocal microscopy of a monoclonal stably incorporated GFP-MYC cell line compared to a MYC RTA circuit cell line, showing treatments with DMSO and with 100 nM dTAG$^{Vm}$. **B.** High throughput microscopy quantification of the GFP-MYC line GFP fluorescence as a function of dTAG$^{Vm}$ treatment dose (normalized to DMSO and 1 µM dTAG$^{Vm}$ (max and min, respectively). Z-factors[3] for the assay were <0 for all treatment doses. **C.** Z′ values[3] for the MYC cell line at all doses with a >0 Z-factor value. **D.** A random small molecule library (3,840) was tested in triplicate on independent weeks against the MYC/STAT3 pool, standard error variance of %GFP+ cells, %mKate2+ cells, and viability (total nuclei count) between the three replicates is shown for all molecules. **E.** All results from the 7,479-molecule library sourced from multiple commercial clinical collections tested against MYC, STAT3, FOXP3, and SHP2 RTA cells. Molecules that showed >5% activity in another circuit are filtered in gray. **F.** Displaying nontoxic (>60% viability) molecules that activated >1 RTA line (multi-hitters, filtered out from analysis in Figure 7). Known or clearly labeled fluorescent compounds that increased GFP or mKate2 fluorescent signal are highlighted in blue (above). (Below): BET inhibiting molecules show robust nonspecific activation of multiple circuits, and HDAC inhibitors seem to show moderate nonspecific activation. **G.** MYC RTA circuits were treated with 10 µM tacrolimus, pimecrolimus, and ascomycin for 24 h and percent GFP+ cells were detected by fluorescence (results were normalized to max (dTAG$^{Vm}$) and min (DMSO) signal. **H.** All results from the CRBN glue library screen (showing extended axis for CK1a), numbers of molecules filtered out for each screen (for appearing as >10% Activity hit in another circuit) are displayed in the legend.

#### Materials and Methods

Data reproducibility

Immunoblot results are representative of n=2 cell treatments performed on independent days (or n=3 when quantified). All flow cytometry results are quantified from n=3 replicates treated and analyzed on independent days. Confocal microscopy images are representative of n≥3 frames from n=2 imaging experiments on independent days. For barcode experiments, replicates were treated and frozen on n=3 independent days, then processed in parallel for RT-PCR and PCR amplification for sequencing. qPCR was conducted similarly, with samples from n=3 sequential passages generated and frozen on separate days prior to parallel downstream processing; qPCR was performed in triplicate technical replicates for each independent replicate. Reporting of confluence for induction of cell death (Figures 5A-C and S5a) and cell survival (Figures 5D-F and S5b) were from n=3 experiments performed on independent days.

For RTA circuit activity by live cell fluorescence or confocal microscopy, n=3 independent and no technical replicates were performed except in two experiments where technical replicates were required to calculate Z-scores: Experiment (1) Comparing endogenously-tagged Acr RTA circuits to HiBit, n=2 technical replicates were included for each of n=3 experiments treated on independent days (Figures 6J-L and S6C), Experiment (2) Generating a Z score for the MYC RTA circuit. For each experimental replicate dose treatment, 24 technical replicates were collected for both GFP-MYC and MYC/STAT3 RTA circuits (Figure S7b: GFP-MYC and Figure 7D and S7c: MYC RTA circuits).

HibiT luminescence dose curves were treated and analyzed by plate reader on three separate days, yielding n=3 independent experiments, each with n=2 technical replicates on the plate. This was compared directly to RTA circuits and direct GFP fusions, plated and analyzed by Image Xpress live cell confocal microscopy on n=3 independent days with n=2 technical replicates per plate, although direct GFP fusions did not have quantifiable signal via this method.

For library screening, variability in fluorescence and viability between n=3 replicates treated and imaged on independent days was assessed using the Stanford SPECS molecule library (Figure S7d). Screens with the clinical collection and CRBN-glue oriented libraries were performed with n=1 replicate.

Statistical analysis

When comparing between two samples, unpaired two-tailed t tests were used; the *P* value was adjusted for multiplicity if multiple t tests were performed within one experiment. An ordinary one-way ANOVA was used to compare between >2 samples; with Dunnett's or Tukey's multiple comparison tests used when samples were compared to control, or to each other, respectively. All replicate data points are shown when possible without detracting from visualization of the data. Analyses were performed in GraphPad Prism, and resultant *P* values are either precisely displayed, or are shown in the supplementary source data file and the following symbols are reported in the main text: *$P<0.05$, **$P<0.01$, ***$P<0.001$, ****$P<0.0001$, and ns ($P>0.05$).

Plasmid construction

The Sleeping Beauty transposase system[4] was used to stably integrate all exogenous genes. The VPR-BFP-*Spy*dCas9 and AcrIIA4 sequences (gifts from Stanley Qi at Stanford University) were cloned into a sleeping beauty backbone using PCR amplification with IDT primers followed by HiFi DNA assembly. Target protein constructs were cloned into the Acr-fusion construct by HiFi DNA assembly on NotI/PacI restriction-enzyme-digested backbone. Many target construct sequences were provided by individual labs through Addgene and are gratefully acknowledged here: MYC provided by Jialiang Wang (Addgene plasmid #46970), PBRM1 provided by William Kaelin (Addgene plasmid #107406), STAT3 provided by Geert van den Bogaart (Addgene plasmid #111934), SMARCA4 provided by Luca Tiberi (Addgene plasmid #153950), the *Sau*dCas9 sequence was provided by Ronald Cohn (Addgene plasmid # 135338), SHP1 and PTPN1 were provided by Ben Neel (Addgene plasmids #8572 and #860), SHP2 was provided by Anton Bennett (Addgene plasmid #12283). BFP was omitted from dCas9A in subsequent constructs due to undetectable expression; all-in-one constructs (Figure 2E) and constructs driving endogenous RTA circuit tagging were engineered using the SpydCas9-VPH CRISPRa backbone (Cellecta). Other sequences were synthesized as gene fragments by Twist Bioscience.

Mammalian cell culture

HEK293T cell lines purchased from the ATCC were a gift from Carolyn Bertozzi at Stanford University. Monoclonal HEK293T cell lines expressing TRE-promoted GFP were a gift from Luke Gilbert at UCSF. Cells were cultured in DMEM media containing 10% heat-inactivated FBS, 100 U/mL penicillin, and 100 µg/mL streptomycin at 37 °C under 5% $CO_2$. Normal passaging procedure was one PBS wash, followed by 2-5 min tryspin incubation at 37 °C, quenching with complete media, and dilution onto a new culture or experimental plate. The jetPRIME transfection kit was used to transfect all DNA and siRNA into cells. For stable transfection, the sleeping beauty transposase (SB100) plasmid was included in the jetPRIME transfection protocol at a 1:10 ratio. Once genes were incorporated, selection pressure was constantly maintained in normal media with one or more of: hygromycin (250 µg/mL), puromycin (2 µg/mL), blasticidin (10 µg/mL), or zeocin (400 µg/mL).

Immunoblot

Plated cells were washed in PBS and lysed in cold RIPA lysis buffer containing protease inhibitor cocktail and 1:1000x diluted benzonase for 20 min on ice, transferred to tubes, and clarified at 12,000 x *g* for 15 min. Concentration was quantified by BCA assay. Proteins were denatured in a final concentration of 1X LDS buffer (NuPAGE) containing 2.5% 2-mercaptoethanol at 95 °C for 5 min. Total protein loading was held constant for a given experiment, and 12-45 µg total protein was separated by gel electrophoresis (4–12% polyacrylamide gel, 180 V, 40-60 min) and transferred to a nitrocellulose membrane. Membranes were blocked at rt for 30-60 min in TBST (PBS+0.1% Tween) containing 5% milk, incubated with primary antibodies overnight at 4 °C in TBST+3% milk, washed 3x 5 min with TBST, incubated at rt with secondary antibody for 1 h in Intercept blocking buffer (LI-COR), and washed 3 more times prior to signal detection with the LICOR Odyssey CLx instrument.

<u>Flow Cytometry</u>

Cells were seeded in a 24-well plate. The next day, cell media was replaced with media containing DMSO or compound treatment (0.1% final DMSO concentration in all samples). 1-2 days after treatment, cells were lifted with trypsin, pelleted by centrifugation (300 x *g*) and analyzed by flow cytometry using either the BD Accuri C6 plus (Figures S3E and S4A) or the BD FACS Symphony A5 flow cytometer (all other flow cytometry figures). Cells were gated in FloJo software by forward and side scatter (FSC-A/SSC-A), followed by single cell gating using (FSC-A/FSC-H), and often for % positive for a fluorescent reporter. Compensation was performed in FloJo software. Flow cytometry histograms are represented on a biexponential axis to show large fluorescence changes or a log axis to demonstrate smaller fluorescence variations.

<u>Live cell fluorescence microscopy (IncuCyte)</u>

The IncuCyte instrument resides within a tissue culture incubator maintained at 37 °C with 5% $CO_2$. Data were acquired using a 10x objective lens, and phase-contrast, (1.24 µm/pixel resolution), GFP fluorescence (excitation: 441-481, emission: 503-544, acquisition time = 300 ms) and mKate2 fluorescence (excitation: 567-607, emission: 622-704, acquisition time = 400 ms) were acquired at predetermined timepoints. All values output were automatically averaged by the instrument from n=2 images within each well. Data were analyzed from the GFP channel alone (integrated GFP intensity, reported as fold-change over control) or from both GFP and phase channels (GFP-positive area as a fraction of total cell area)

Briefly, cells were lifted and plated at 10,000 cells/well in 90 µL of complete media in 96 well flat-bottom plates and placed in a humidified incubator overnight. The next day, cells were treated by adding 10 µL of 10X concentrated degrader molecules or DMSO control in media (final 1X concentration in 0.1% DMSO). Alternatively, 10 µL jetPRIME DNA or siRNA transfection mixtures were added. Cells were placed in the Incucyte instrument and scanned at the timepoints noted in each figure legend. Image analysis was empirically optimized for these circuits and fluorophores using the associated IncuCyte live cell imaging and analysis software. For quantifying confluence/cell area, the "Classic Confluence" package was used with a 0.8 segmentation adjustment and a minimum area of 200 $\mu m^2$ to gate-out debris. GFP and mKate2 were quantified using the "Surface fit" segmentation module, threshold = 2 GCU (for GFP) 0.4 GCU (for mKate2). *Sau*dCas9-containing circuits had lower average fluorescence and GFP threshold was reduced to 1.2 GCU in those experiments (Figure 4).

For cell survival and cell death inducing circuits (Figure 5), 4,000 cells in 90 µL were plated per well of a 96 well flat-bottom plate and allowed to adhere overnight. The following day various $dTAG^{Vm}$ concentrations or DMSO was added in 10 µL of normal media, and phase image was quantified every 4 hours for 72 hours.

Endogenous tagging

Endogenous tagging was performed using the PITCh system.[5] Briefly, HEK293T cells in a 6-well dish were transiently transfected following the jetPRIME protocol with 1.2 µg of plasmid that contains target sgRNA, PITCh sgRNA, and *Spy*Cas9, and 0.6 µg of donor vector (5'HA-blast-2A-TAG-Flag-$FKBP12^{F36V}$-3'HA, where HA are the homology regions N-terminal to the endogenous knock in site, and TAG = Acr, eGFP, mNG11, or HibiT). Cells were split after 1 day onto a 10 $cm^2$ dish. Three days after transfection, cells were selected with blasticidin (10 µg/mL). After 2 weeks and complete death of control cells, constructs containing *Spy*dCas9-VPH, LgBit, or mNG1-10 were stably integrated with Sleeping beauty transposase and cells were selected with hygromycin (50 µg/mL) for 2 weeks. For Acr lines, cells were treated with 1 and 10 nM of $dTAG^{Vm}$ for 24 h, and monoclonal cells were sorted for GFP+ to achieve the monoclonal endogenously tagged circuit line. Endogenously tagged eGFP and mNG -fusions were sorted for green fluorescence, and HibiT lines were sorted only on single cells (gating in Figure S6A). Endogenous tagging was confirmed by immunoblot using anti-FLAG and target-specific antibodies to assess knock-in efficiency and loss of wild-type protein.

| **Endogenous Target** | **sgRNA sequence** | **5' HA** | **3' HA** |
|---|---|---|---|
| MEK1 | GATGGGCGTCGGCTTCTTCT | TACCCGGGTCCAAAATGCCCAAG | AAGAAGCCGACGCCCATCCAGCTG |
| PARP1[6] | GGGGAGGATGGCGGAGTCTT | GCTAGGGGAGGATGGCGGAGTCT | TCTTCGGATAAGCTCTATCGAGT |
| RIOK3[6] | CCTTCATTCCCGAATGGATC | GTCTCTGCCTTCATTCCCGAATG | ATGGATCTGGTAGGAGTGGCATC |
| PRMT9[6] | GCCTGGGCCGCGAGTTCGACA | CCATACAAGTGGTGACTGCCATG | ATGTCGAACTCGCGGCCCAGGTC |
| ATM[7] | GATCATTAAGTACTAGACTCA | CCTATATGTATTTTTTTACAGACAGTGATGTGTGTTCTGAAATTGTGAACC | CTAGTACTTAATGATCTGCTTATCTGCTGCCGTCAACTAGAACATGATAGAG |

Immunofluorescence microscopy

Cells were plated at 35,000 cells/well in 450 µL an 8-well chamber slide pre-coated with 10 µg/mL fibronectin. The next day, cells were treated with 50 µL solutions of media containing DMSO or 10X molecule degrader (final cell treatment: 1 µM in 0.1% DMSO). After the indicated length of time in the figure legend, cells were fixed with 4% formaldehyde in PBS for 10 min, dark. Cells were washed 2x by media replacement with PBS and permeabilized with 0.3% Triton-X-100 in PBS, 10-15 min room temperature. Blocking was performed for 1 hour at room temp with 3% Goat serum in PBS, then the media was replaced with primary antibody in 3% Goat serum in PBS and stained overnight at 4 °C. The next day, cells were washed 2x 5 minutes each and media was replaced with secondary antibody in 3% Goat serum in PBS for 1 hour at room temperature. Cells were washed 3x in PBS 5 minutes each, 1 µg/mL DAPI stain was included in the first wash. Cells were imaged with a Nikon A1R confocal microscope using a Plan Fluor 60X oil immersion, 1.30-numerical aperture objective. The microscope was equipped with a 405-nm violet laser, a 488-nm blue laser, a 561-nm green laser, and a 639-nm laser. Images were analyzed with FIJI software.

Barcode sequencing

The 6 cell lines with unique barcode reporters were lifted, counted, and pooled to an even mixture. On a 6-well plate, 400,000 total cells/well were plated in 2 mL media. The following day, media was replaced with media containing 0.1% DMSO or treatment compound. After 24 hours, cells were lifted with trypsin, quenched with media, transferred to centrifugation tubes, pelleted at 300 x *g*, and washed 1x with 1 mL PBS. Cells were pelleted again and frozen at -80 °C until the next step. RNA was extracted using the RNeasy mini kit. Extracted total RNA was converted to cDNA using 1 µg template RNA and the TaqMan reverse transcription kit at 25 °C for 10 min, 37 °C for 2 h, and 95 °C for 5 min, using 2.5 µM oligodT. Finally, cDNA was amplified using 2x AmpliTaq Gold mix from 2 µL of template cDNA Forward primer: CAACGTCAAGATCAACGGGG, Reverse primer: CTCCCCCTGAACCTGAAACAT; Cycle number (20–25) was determined empirically by qPCR prior to amplification. Unpurified PCR reactions were submitted to Quintara Biosciences for amplicon sequencing by nanopore reporting full length fastq reads. Only exact-match barcode sequences were counted, and abundance relative to DMSO controls reported.

qPCR

One million cells were collected from each cell line per replicate on independent passages. Cells were lifted with trypsin, quenched with normal media, pelleted at 300 x *g* for 5 min, washed with 1 mL PBS, and pelleted again. Cell pellets were stored at -80 °C. RNA was extracted using the RNeasy mini kit. Extracted total RNA was converted to cDNA using 1 µg template RNA and the TaqMan reverse transcription kit at 25 °C for 10 min, 37 °C for 2 h, and 95 °C for 5 min, using 2.5 µM hexamers (Figure 2F) or 2.5 µM oligodT (all other qPCR data). For the all-in-one construct, random hexamers were used to mitigate RT drop-off, as the qPCR amplicon is located >6 kb from the poly(A) tail in that construct. qPCR was performed on 2 µL cDNA using a Bio-Rad CFX96/C1000 Touch system with PowerUp™ SYBR™ Green Master Mix (20 µL total volume). Cycling conditions were: UDG activation 50 °C 2 min; polymerase activation 95 °C 2 min; 40 cycles of 95 °C 15 s, 60 °C 1 min. The following primers were used at 500 nM final concentration:

GAPDH primers[8] FP: AAACAGCAGATTCGCCTGGA, RP: TCATCCGCTCGATGAAGCTC
Cas primers[8] FP: TCCAAAATCAAGTGGGGCGA, RP: TGATGACCCTTTTGGCTCCC
Acr primers: FP: GACGGCAACGAGTACGTGAT, RP: ACTCCTGGTTCCAACCGTTC

Cq values were determined by Bio-Rad CFX Maestro software, and ΔCq was calculated by subtracting GAPDH Cq from target Cq. Relative transcript levels were determined using $2^{-\Delta Cq}$.

High-throughput chemical screening

For the random compound pilot library, the first 3,840 of the Stanford SPECS chemical library (a diversified/randomized library of Lipinski-compliant molecules) were chosen. First, cMYC and STAT3-sensing cell lines were pooled together 1:1 and 30 µL cells in complete media were seeded into 384-well black-sided clear-bottom plates using a WellMate Microplate Reagent Dispenser with integrated stacker to a final seeding density of 2,500 total cells/well. Cells were allowed to settle 45 min before transferred to an incubator (37 °C, 5% $CO_2$). The next day, 60 nL of 5mM stock compounds were dispensed to columns 3-22 acoustically with Echo 655 liquid handler and Spinnaker Robot System to final 10 µM concentration in each well. Positive control (100 nM final concentration $dTAG^{Vm}$) was added to columns 1-2. One day after compound addition, 10 µL of 10 µM (4x) Hoechst dye was added in media using the Microplate Reagent dispenser to a final 2.5 µM concentration. After 30 min, plates were automatically imaged for Hoechst, GFP, and mKate2 fluorescence using a ImageXpress Micro Confocal paired with an automated microplate handler (PF400), a Cytomat $CO_2$ incubator, a Microscan barcode reader, and the AWES Green Button Go Automated Scheduler software. Imaging took ~15 min/plate. Image analysis was performed with the MetaXpress software using the multi-wavelength cell scoring protocol with gating widths: (7- 50 µm) and gating intensity above local background 250 graylevels (GFP, Hoechst), and 200 graylevels (mKate2). Data were gathered in triplicate on independent days, and for each plate %GFP+ and %mKate2+ cells were normalized to max and min (median fluorescence in $dTAG^{Vm}$-treated wells and median plate fluorescence, respectively). Variance between replicates is reported.

The clinical compound library used in Figures 7H and 7I was assembled from five commercial sources (Biomol, Microsource, Selleckchem, LOPAC, and MedChemExpress), resulting in a collection containing both unique compounds and inter-library duplicates. The library comprised 7,479 compounds, of which 131 were excluded as known fluorescence assay interferents, leaving 7,348 for analysis. Screening was perfumed as described above, with the MYC/STAT3 pool screened on one day, and SHP2/FOXP3 lines pooled and screened on another. Percent viability was calculated from Hoescht nuclei counting compared to median plate cell counts. %GFP+ and %mKate2+ cells were normalized to max and min (median fluorescence in dTAG$^{Vm}$-treated wells and median plate fluorescence, respectively). Filtering: for labeling molecules as specific or nonspecific, any molecule with >5% activity against >1 cell line were filtered (all data points shown in Figure S7).

The CRBN molecular glue library was assembled in the laboratory of Nathanael Gray comprising molecules from multiple degrader discovery campaigns. Screening was performed as described above, with the addition of 100 nM dTAG-48 alongside dTAG$^{Vm}$ as a positive control. The best-performing degrader in the screen was used for maximum normalization. As some compounds exceeded positive control activity, values above 100% are observed for this library. Filtering was performed on >10% activity against >1 cell line.

<u>Comparing live-cell HiBiT detection, eGFP, and RTA circuits (Figure 6J-K).</u>

Cells were plated at 5,000 cells/well in 30 µL into 384-well black-sided clear-bottom plates using a multichannel pipet. GFP and RTA cell lines were plated in complete normal media; HiBit cell lines were plated in cOptiMEM (OptiMEM with GlutaMAX, 5% FBS, 1% MEM non essential amino acids solution (Gibco), and 1mM sodium pyruvate). White-sided plates were avoided for HiBiT assays due to significant well-to-well signal bleed-through in 384-well format. The next day, 10 µL of 10x concentrated dTAG$^{Vm}$ treatments or DMSO controls in complete media were added by multichannel (final 0.1% DMSO in all wells). After 24 hours, Nano-Glo® Live Cell Substrate was diluted 20x in LCS dilution buffer according to the Promega protocol and 10 µL was added to HiBiT assay wells. Wells were shaken for approximately 1 min prior to luminescence detection on a SpectraMax iD5 plate reader (Molecular Devices). For GFP and RTA circuits, 10 µL of 10 µM Hoechst dye in complete media was added to the wells, plates were left in the incubator for 30+ mins of incubation with Hoechst prior to imaging the plates for Hoechst, GFP, and mKate2 fluorescence using the ImageXpress Micro Confocal.

<u>CRBN-dependency by RTA circuit assay</u>

Molecular glue lead molecules from the screen were assessed for RTA circuit activation and CRBN independence by live cell fluorescence microscopy (IncuCyte). Briefly, 10,000 UBE2S RTA cells were plated in a 96 well flat-bottom plate. The next day, wells were pre-treated for 2 hours with either DMSO control or CRBN-6-5-5-VHL to deplete CRBN E3 from cells, before molecules at were added to a final concentration of 10 µM and cells were assessed for mKate2+ fluorescence /phase area by IncuCyte.

#### Modeling RTA circuits

Detailed description of the ODE model including parameters used to generate Figures 2A, 2B, and 2I, are appended to the end of this supplementary file

#### Equipment

| Name | Manufacturer | Catalog No. |
|---|---|---|
| WellMate Microplate Reagent Dispenser | Matrix | 201-10001 |
| WellMate Microplate Stacker | Matrix | 201-20001 |
| Echo 655 liquid handler | Beckman Coulter | 001-16080 |
| ImageXpress Micro Confocal High-Content Imaging System | Molecular Devices | |
| Spinnaker Robot System | Thermo Scientific | F01981 |
| Precise Automation PF400 Microplate Handler | | |
| Cytomat 2C automated $CO_2$ incubator | Thermo Fisher, | |
| Microscan Barcode Reader | Molecular Devices | SR-710 |
| Trans-Blot Turbo Transfer System | BioRad | 1704150 |
| Li-Cor Odyssey CLx Imaging System | LI-COR | |
| IncuCyte S3 Live-Cell Analysis Instrument | Sartorius | |
| BD Accuri C6 plus | BD Biosciences | |
| BD FACS Symphony A5 | BD Biosciences | |

#### Software

| Name | Manufacturer | Version or Cat. No |
|---|---|---|
| AWES Green Button Go Automated Scheduler software | Molecular Devices | PF00-MA-00400 |
| MetaXpress High-Content Image Acquisition and Analysis Software | Molecular devices | Version 6.7 |
| GraphPad PRISM | Dotmatics | Version 10 |
| Adobe illustrator | Adobe | Yr 2024 |
| Microsoft office | Microsoft | 16.83 |
| FIJI | Schindelin *et al* [9] | Version 2 |
| Python | Open source | |

#### General laboratory materials

| Name | Manufacturer | Catalog No. |
|---|---|---|
| HEK293T cells | ATCC | CRL-3216 ™ |
| 384-well black-sided clear-bottom plates | Corning | 3764 |
| DMEM +L-Glut, High Glucose | Fisher (Gibco) | 31053028 |
| FBS (heat inactivated) | Fisher Scientific | 10-438-026 |
| Pen-Strept, 100X liquid | Fisher (Cytiva HyClone) | SV30010 |

| | | |
|---|---|---|
| Corning™ Cell Culture Phosphate Buffered Saline (1X) | Fisher (Corning) | MT21030CV |
| Hoechst 33342 | Thermo | 62249 |
| 0.25% Trypsin + 2.21 mM EDTA 1x [-] sodium bicarbonate | Fisher Scientific | SH3004201 |
| dTAG$^{V}$ | Synthesized in house | CAS: 2624313-15-9 |
| RIPA Lysis buffer | Thermo Fisher Scientific | 89901 |
| Pierce BCA Protein Assay Kit, | Thermo Fisher Scientific | 23227 |
| NuPAGE LDS Sample Buffer (4x) | Thermo Fisher Scientific | NP00007 |
| 2-Mercaptoethanol | Sigma Aldrich | 63689 |
| 4–12% Criterion™ XT Bis-Tris Protein Gel, 18 well, 30 µl | BioRad | 3450123 |
| Trans-Blot Turbo RTA Midi 0.2 µm Nitrocellulose Transfer Kit | BioRad | 1704271 |
| Corning Cell Culture Phosphate Buffered Saline (1x) | Thermo Fisher | MT21040CV |
| NotI-HF | NEB | R3189L |
| PacI | NEB | R0547L |
| Q5 High-Fidelity 2X Master Mix | NEB | M0492L |
| NEBuilder HiFi DNA Assembly Master Mix | NEB | E2621S |
| Nunc™ Lab-TEK™ II Chambered Coverglass | Fisher Scientific | 12-565-338 |
| Human Plasma Fibronectin purified protein | Millipore Sigma | FC010-1mg |
| Gibco Goat Serum, New Zealand origin | Fisher Scientific | 16-210-072 |
| Methanol-free Formaldehyde Ampules, Thermo Scientific, 15% Formaldehyde (w/v) | Fisher Scientific | PI28906 |
| Triton-X-100, for molecular biology | Sigma | T8787-50mL |
| RNeasy Mini Kit | Qiagen | 74104 |
| Qiaquick PCR purification kit | Qiagen | 28104 |
| AmpliTaq Gold 360 Master mix | Thermo Fisher | 4398881 |
| TaqMan Reverse Transcription Reagents | Thermo Fisher | N8080234 |
| Corning Costar 96-well, cell culture-treated Flat bottom microplate | Fisher Scientific | 07-200-89 |
| Intercept® (PBS) Blocking Buffer | LI-COR | NC1660553 |
| Hygromycin B (50 mg/mL) | Thermo Fisher | 10-687-010 |
| Puromycin (10 mg/mL) | Invivogen | 58-58-2 |
| Blasticidin S HCl (10 mg/mL) | Thermo Fisher | A1113903 |
| Zeocin 100 mg/mL in HEPES, sterile filtered | Thermo Fisher | AAJ671408EQ |
| Opti-MEM™ I Reduced Serum Medium, GlutaMAX™ Supplement | Thermo Fisher | 51985034 |
| MEM Non-Essential Amino Acids Solution (100X) Gibco™ | Thermo Fisher | 11140050 |
| Sodium Pyruvate (100 mM) Gibco™ | Thermo Fisher | 11360070 |
| Nano-Glo Live Cell Assay System, 100 assays | Promega | N2011 |
| 6-well cell culture plates | Genessee | 25-105 |
| 24-well cell culture plates | Genessee | 25-107 |

#### Antibodies

| **Name** | **Dilution or concentration** | **Catalog No. & Manufacturer** |
|---|---|---|
| Monoclonal ANTI-FLAG M2 antibody produced in mouse, clone M2 purified immunoglobulin, buffered aqueous solution | 1:500: Fluorescent microscopy<br>1:1000: Immunoblot | F3165-1mg<br>Sigma Aldrich |
| Vinculin Antibody | 1:1000: Immunoblot | #4650 Cell Signaling Tech. |
| β-Actin (13E5) Rabbit mAb | 1:4000 Immunoblot | #4970 Cell Signaling Tech. |
| Goat Anti-Mouse IgG H&L (Alexa Fluor® 488) preadsorbed | 2 µg/mL: Fluorescent microscopy | ab150117, Abcam |
| Goat Anti-Mouse IgG H&L (Alexa Fluor® 568) preadsorbed | 2 µg/mL: Fluorescent microscopy | ab175701, Abcam |
| IRDye® 800CW Goat anti-Mouse IgG (H + L), | 1:10,000: Immunoblot | NC9401841, LI-COR |
| IRDye 680RD Goat anti-Rabbit IgG Secondary Antibody | 1:10,000: Immunoblot | NC025229, LI-COR |
| ATM (D2E2) Rabbit mAb #2873 | 1:1000: Immunoblot | 2873S, CST |
| MEK1 (61B12) Mouse mAb #2352 | 1:1000: Immunoblot | 2352S, CST |
| RIOK3 Polyclonal antibody | 1:1000: Immunoblot | 13593-1-AP, Proteintech |
| PARP1 Polyclonal antibody, 500 ug ml | 1:1000: Immunoblot | 13371-1-AP, Proteintech |
| CK1a (Rb) | 1:1000: Immunoblot | #ab108296, abcam |
| WEE1 (Rb) | 1:1000: Immunoblot | 13084, CST |
| UBE2S (Rb) | 1:1000: Immunoblot | 11878, CST |
| Actin (Ms) | 1:5000: Immunoblot | 3700, CST |
| CRBN (Rb) | 1:1000: Immunoblot | NBP1-91810, Novus |

7. OpenCell: Endogenous tagging for the cartography of human cellular organization | Science https://www.science.org/doi/10.1126/science.abi6983.

8. DNA targeting specificity of RNA-guided Cas9 nucleases | Nature Biotechnology https://www.nature.com/articles/nbt.2647.

9. Schindelin, J., Arganda-Carreras, I., Frise, E., Kaynig, V., Longair, M., Pietzsch, T., Preibisch, S., Rueden, C., Saalfeld, S., Schmid, B., et al. (2012). Fiji - an Open Source platform for biological image analysis. Nat Methods *9*, 10.1038/nmeth.2019. https://doi.org/10.1038/nmeth.2019.

### A simplified model of the relationship between anti-CRISPR, Cas, and the expression of a target gene

Here, GFP ($G$) production is modulated by binding of free Cas ($C$) to the TRE promoter ($T$) upstream of the GFP gene. GFP expression is positively regulated by promoter-bound Cas ($T_b$), acting as an activator in this scenario. Anti-CRISPR (A) binds to Cas to make an Acr-Cas complex ($X$) which competitively inhibits Cas binding to promoter.

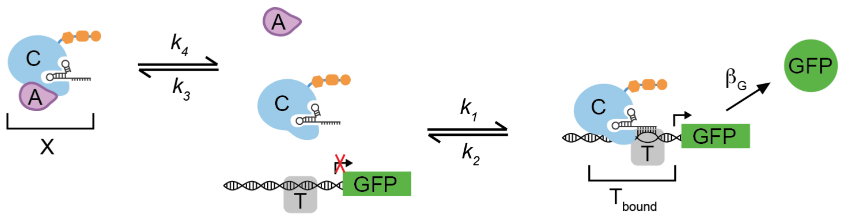


For simplicity, transcription and translation are modeled as a single effective step. Because guideRNA binds Cas with high affinity, we assume all Cas is sgRNA-bound and do not model Cas:guideRNA complex formation. Additionally, several intermediate processes involved in Cas-mediated DNA binding, such as PAM recognition, DNA unwinding, and guideRNA:DNA interactions, which have previously been modeled explicitly *(Farasat and Salis, 2016)*, are condensed into a single effective step representing formation of the Cas:RNA:DNA complex ($T_b$).

We can describe transcription factor binding and gene production using the following relationships:

$$A + C \underset{k_4}{\overset{k_3}{\rightleftharpoons}} X$$

$$C + T \underset{k_2}{\overset{k_1}{\rightleftharpoons}} T_b$$

$$T_b \xrightarrow{\beta_G} T_b + G$$

For each of our proteins, we can also include a synthesis rate constant $(\beta_C, \beta_A)$ as well as natural degradation/dilution constants $(\gamma_C, \gamma_A, \gamma_X,)$

$$Gene_A \xrightarrow{\beta_A} Gene_A + A$$

$$Gene_C \xrightarrow{\beta_C} Gene_C + C$$

$$A \xrightarrow{\gamma_A} \emptyset$$

$$C \xrightarrow{\gamma_C} \emptyset$$

$$X \xrightarrow{\gamma_X} \emptyset$$

$$G \xrightarrow{\gamma_G} \emptyset$$

Finally, we can include our added induced degradation component ($\gamma$), analogous to dTAG treatment. Although the model permits arbitrarily high degradation rates, biological reality imposes constraints as degradation machinery can become rate-limiting, and degrader molecules such as dTAG are subject to a hook effect that is not captured here. Here we consider two RTA circuit possibilities, in Model 1, $\gamma$ degrades Acr from the Cas:Acr complex, and Model 2 is a model of Cas-codegradation, in which Cas:Acr complex is degraded by $\gamma$. These two models represent limiting cases; in practice, the degree of Cas9 co-degradation may be partial

| **Model 1** | **Model 2** |
|---|---|
| $A \xrightarrow{\gamma} \emptyset$ | $A \xrightarrow{\gamma} \emptyset$ |
| $X \xrightarrow{\gamma} C$ | $X \xrightarrow{\gamma} \emptyset$ |

**Model 1** is described by the following system of ordinary differential equations, where $[T] = [T_{tot}] - [T_b]$ (total promoter is conserved), $\theta = [T_b]/[T_{tot}]$, and $K = k_2/k_1$:

$$\frac{d[G]}{dt} = \beta_G \left( \frac{\theta^n}{K^n + \theta^n} \right) - \gamma_G[G] \tag{1}$$

$$\frac{d[T_b]}{dt} = k_1[C][T] - k_2[T_b] \tag{2}$$

$$\frac{d[C]}{dt} = \beta_C + k_2[T_b] + k_4[X] - \gamma_C[C] - k_1[C][T] - k_3[A][C] + \gamma[X] \tag{3}$$

$$\frac{d[A]}{dt} = \beta_A + k_4[X] - [A](\gamma_A + \gamma) - k_3[A][C] \tag{4}$$

$$\frac{d[X]}{dt} = k_3[A][C] - k_4[X] - [X](\gamma_X + \gamma) \tag{5}$$

**Model 2 (Cas-co-degradation): single-equation modification of Model 1**

$$\frac{d[C]}{dt} = \beta_C + k_2[T_b] + k_4[X] - \gamma_C[C] - k_1[C][T] - k_3[A][C] \tag{6}$$

Note that GFP production is governed by a Hill-type activation function of promoter occupancy $\theta = [T_b]/[T_{tot}]$, where $0 \leq \theta \leq 1$, allowing changes in $C$ to modulate promoter binding without enforcing a strictly linear relationship with gene expression. The codegradation of Cas model (Model 2) replaces equation (3) with equation (6).

Model 1 and 2 equations were numerically solved using Python's solve ivp function from scipy.integrate. Parameters and initial starting concentrations are described in Tables 1–3. The initial conditions are set to 0 for all species. Total DNA ($[T_{tot}]$) is held constant as a fixed parameter ($[T_{tot}] = 0.1$). Only Model 1 is reported in Figures 2A and B, Figure 2I compares both models. Although parameters are arbitrary and unitless, they reflect a biologically motivated hierarchy in which binding is favored over unbinding. Key qualitative behaviors were found to be robust across a wide range of parameter values.

| **Parameters** | **What the parameters are set to for Fig 2A/B and Fig 2I** |
|---|---|
| Cellular synthesis rates ($\beta_G$, $\beta_C$) | 10 |
| Cellular synthesis rate of anti-CRISPR ($\beta_A$) | (Fig 2A/B): varied from $10^0$-$10^4$ (Fig 2I): 100 |
| Cellular degradation rates ($\gamma_C$, $\gamma_A$, $\gamma_G$) | 1 |
| Induced degradation rate ($\gamma$) | (Figs 2A/B and 2I): varied from $10^{-3}$ to $10^3$<br>Fig 2I (timecourse): 0 at t=0, then 10 |
| Binding rates ($k_1$, $k_3$) | 100 |
| Unbinding rates ($k_2$, $k_4$) | 10 |
| Total DNA concentration ($T_{tot}$) | 0.1 |
| Hill coefficient ($n$) | 2 |

Table 1: Parameter values used in the model. Parameters $\beta_A$ and $\gamma_{deg}$ were varied across the ranges shown for certain figures; all other parameters were fixed.

| **Dependent variables** | **Initial conditions** |
|---|---|
| Gene expression level ($G$) | 0 |
| Concentration of dCasA ($C$) | 0 |
| Concentration of Acr ($A$) | 0 |
| Concentration of dCasA:ACR complex ($X$) | 0 |
| Concentration of bound transcription factor ($T_b$) | 0 |

Table 2: Initial conditions for dependent variables for steady state approximation.

| **Independent variables** | **Values for steady state approximation** |
|---|---|
| Time ($t$) | 1000 |

Table 3: Independent variable.

For steady-state approximations (Figures 2A, 2B, and the right panel of 2I), t=1000 was used to approximate steady state. For the degradation timecourse (Figure 2I, ACR and Cas degradation panel), the system was first solved to steady state with no induced degradation ($\gamma = 0$); the resulting dependent variable concentrations at steady state were then used as initial conditions for a second simulation with all parameters held constant except $\gamma = 10$
